# Disease and introgression explain the long-lasting contact zone of Modern Humans and Neanderthals and its eventual destabilization

**DOI:** 10.1101/495515

**Authors:** Gili Greenbaum, Wayne M. Getz, Noah A. Rosenberg, Marcus W. Feldman, Erella Hovers, Oren Kolodny

## Abstract

Neanderthals and modern humans both occupied the Levant for tens of thousands of years prior to modern humans’ spread into the rest of Eurasia and their replacement of the Neanderthals. That the inter-species boundary remained geographically localized for so long is a puzzle, particularly in light of the rapidity of its subsequent movement. We propose that disease dynamics can explain the localization and persistence of the inter-species boundary; in this view, each species carried pathogens to which it was largely immune and tolerant, but that could spread to the other, vulnerable, species, inducing a significant disease burden. Epidemics and endemic diseases along the interspecies boundary would have mitigated against bands of one species migrating into regions dominated by the other species. Together with decreased population densities and limited inter-group interactions due to disease burden, this mechanism could have resulted in a fixed and narrow contact-zone. We further propose, and support with results from dynamical systems models, that genetic introgression, including transmission of alleles related to the immune system, would have gradually allowed the two species to overcome this barrier to pervasive inter-species interaction, leading to the eventual release of the inter-species boundary from its geographic localization. Asymmetries between the two species in the initial size of their associated “pathogen package” could have created feedback loops that influenced the rates at which immunity to and tolerance of the novel pathogens were acquired. These asymmetries could have allowed modern humans to overcome the disease burden earlier than Neanderthals, giving them a significant advantage in their subsequent spread into Eurasia, particularly upon interaction with Neanderthal populations that had previously been far from the original contact zone in the Levant.

## 1 Introduction

The lineages leading to modern humans (*Homo sapiens*; henceforth “Moderns”) and Neanderthals (*H. neanderthalensis*) diverged 500-800kya, with Neanderthals inhabiting Eurasia and Moderns inhabiting Africa [1–3]. Migrating out of Africa, Moderns reached the Levant tens of thousands of years prior to their further spread throughout Eurasia, whereas Neanderthals seem never to have spread south of the Levant [4–8]. The two species most likely interacted in this region, at least intermittently, for extended periods of time, along a fairly narrow front [5, 9–13]. This contact is generally believed to have been accompanied by a low level of repeated interbreeding [12–18]. Later, around 45-50kya, Moderns spread further into Eurasia, replacing the Neanderthals within a few thousand years, by about 39kya [5, 6, 11, 19–27] (Fig. 1). This replacement has been extensively studied and debated (e.g. [28–37]). Less attention, however, has been given to the fact that contact in the Levant was made much earlier than the initiation of the replacement phase [11]. Therefore, an important question remains open: Regardless of the driving mechanisms of the eventual replacement, what took so long for replacement to commence? This question is especially puzzling because the time frame in which the interaction front was geographically confined to the Levant [5, 11, 38] was much longer than the time frame in which replacement occurred across Eurasia.

**Figure 1:**
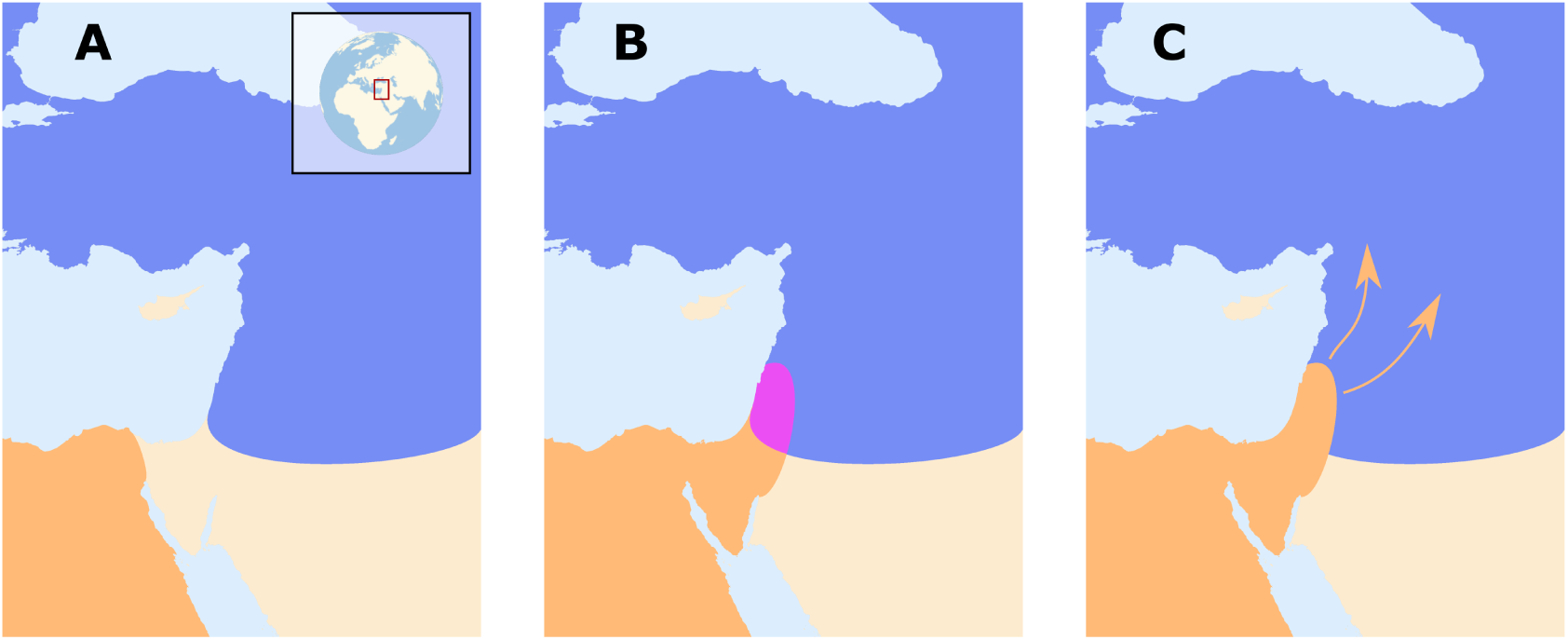
Schematic representation of Modern Human migration and interaction with Neanderthals. The geographic range of Modern humans appears in orange, that of Neanderthals is in blue, and the contact zone in the Levant is in purple. (A) Neanderthals and Moderns were separated for several hundred thousand years, with Moderns co-evolving with tropical pathogens in Africa and Neanderthals co-evolving with temperate pathogens in Eurasia. (B) As early as 180120kya, Moderns began migrating out of Africa into the Levant. Their range remained restricted to this region for tens of thousands of years, during which they interacted intermittently with Neanderthals. We propose that during this period, each species was exposed to novel pathogen packages carried by the other species and experienced disease burden. (C) Around 45–50kya, the inter-species dynamics destabilized and Moderns began expanding to Eurasia. Within several thousand years, Moderns replaced Neanderthals throughout Eurasia.

In this study, we propose that disease dynamics provide a possible solution to this puzzle. We suggest that, following contact, transmissible diseases might have played a prominent role in stabilizing and localizing the inter-species population dynamics in the Levant. We also propose that these dynamics might have resulted in transmissible diseases playing an important role during the later replacement phase. After several hundred thousand years of independent evolution, Neanderthals and Moderns likely acquired immunity and tolerance to different suites of pathogens — a temperate pathogen package in the case of Neanderthals and a tropical pathogen package in Moderns. The re-establishment of contact in the Levant would have resulted in exposure of each species to novel pathogens carried by the other species, which could have spread to the new susceptible hosts, placing a considerable disease burden on both species.

Whether the two groups constitute two species or sub-species is a matter of debate [39–42]; for our purposes, all that matters is that they were geographically distinct for a long period of time, and we refer to them as species for convenience. Nevertheless, many genomic studies of Neanderthals and Moderns have detected a signature of introgression (gene flow) between the species, to the extent that Neanderthal sequences represent 1%-3% of present-day non-African Modern human genomes [15, 16, 43]. That inter-species contact was potentially sufficient to allow for gene flow suggests that disease transmission between the species was likely. Although many of the pathogens that might have been transmitted might not exist today, several genomic studies record potential signatures of events in which pathogens were transmitted between Moderns and Neanderthals, or Moderns and other archaic humans [44–48]. Moreover, several studies have identified signatures of positive selection on putatively introgressed Neanderthal genes in Moderns, particularly in genomic regions such as the MHC complex that are associated with the immune system [2, 43, 47, 49–53], which suggests that disease burden due to inter-species pathogen transmission was significant.

We use mathematical models of inter-species interaction to explore the possible ecological and demographic consequences of exposure to novel pathogen packages upon inter-species contact, coupled with immune-related adaptive introgression. Due to the disease burden imposed by contact with the other species, bands of individuals in either species would have been strongly disadvantaged when migrating into regions dominated by the other, because such migrations would have resulted in increased exposure to novel pathogens. Additionally, disease burden — realized as recurring epidemics, greater endemic pathogen load, or both — would have decreased population densities of both species near the contact front, further reducing the likelihood and motivation for bands of one species to migrate into the range of the other species. The interaction between the two species would thus have been limited to a circumscribed region, which would have been geographically localized by the disease dynamics (Fig. 1). Some contact between the two species would have continued along this front, resulting in pathogen spillovers between the species [54], but also in occasional interbreeding through which transmission of immune-related genes would have occurred. Under these circumstances, bands of each species close to the interaction front would gradually have been able to adapt and acquire tolerance to the novel pathogens, through adaptive introgression as well as by selection on de novo immune-related mutations. Eventually, this process would have reduced disease burden, diminishing the effect of disease transmission dynamics and allowing other processes to drive population dynamics. At this point, the barrier to full inter-species contact and cross-regional migration would have been removed, destabilizing the front of interaction and paving the way for the species dynamics that eventually led to Neanderthal replacement. Disease dynamics, as we will show, are sufficient to explain the extended existence of a stationary interaction front in the Levant, although our analysis does not preclude the importance of additional processes, such as adaptation of the two species to their respective local environments and inter-species competition (e.g. [55–57]; see Discussion).

Once the interaction front was destabilized, presumably around 45–50kya, other processes, previously overshadowed by disease burden, would then have been responsible for the replacement of Neanderthals by Moderns [34, 36, 37, 58], although it has been suggested that disease transmission dynamics could also have played a prominent role in the replacement process [53, 59, 60]. Our model supports this possibility: because conditions relating to disease dynamics need not have been symmetric between Moderns and Neanderthals — for example, the Moderns’ tropical pathogen package may have been more burdensome to the Neanderthals than the Neanderthals’ temperate pathogen package was to Moderns, following the pattern of decreasing pathogen burden with latitude [61, 62] — the Moderns may have overcome the disease burden from contact sooner than Neanderthals. This would have allowed bands of Moderns to migrate into the Neanderthal regions unhindered by novel transmissible diseases, while carrying contagious diseases to which the Neanderthals were not yet immune. Moreover, after the historical front of interaction was crossed and migration reached deeper into Eurasia, this relative Modern advantage would have increased further, as Neaderthal bands encountered far from the intial contact zone would have been intolerant to the entirety of the novel pathogen package spread by the Moderns. We thus suggest, following patterns that occurred multiple times in the colonial era when two long-separated populations renewed contact [48, 63–71], that the replacement of Neanderthals by Moderns may have been facilitated by pathogens to which Moderns were largely immune but to which the Neanderthals were vulnerable.

## 2 Results

Our main proposition is that a persistent Modern–Neanderthal front of interaction in the Levant can be explained by disease burden that prevented each species from expanding to the region dominated by the other. We propose that this front could eventually have collapsed due to immune-related adaptive introgression. In order to (i) demonstrate the feasibility of this scenario, (ii) understand the consequences of variation in the details of this process, (iii) investigate the impact of feedback between disease and gene transmission, and (iv) explore the robustness of our proposition, we model disease transmission and introgression dynamics between two species using dynamical-systems models.

We first explore a model that bridges two independently-treated time-scales, ecological and evolutionary (“two-time-scales model”), as summarized in Figure 2. At the ecological time scale, disease spillovers of novel pathogens, whose impact is measured as the proportion of the non-tolerant population that is infected by each epidemic, are modeled in terms of between- and within-species contact rates (denoted *β*_*ij*_, the contact rates from species *i* to species *j*), using a well-mixed SIR modeling framework (Eq. 2). On the evolutionary time scale, the two species respond to disease burden (*D*_*i*_ for species *i*), which we measure as proportional both to the number (or diversity) of novel pathogens to which the species is exposed (*P*_*i*_) and to the impact of a pathogen at the ecological time-scale (*F*_*i*_); Eq. 1. At each time step in the evolutionary time-scale, this response is modeled as an adjustment of the contact rates (*β*_*ij*_), in proportion to the disease burden experienced (Eq. 4); for example, groups may adjust their tendency to accept individuals from other groups from the same or from different species in response to the impact of disease that they experience. Additionally, also at the evolutionary time-scale, through inter-species contact, immune-related genes are exchanged, decreasing the potential impact of transmissible diseases over time. We assume that the rate of this adaptive introgression is proportional to the inter-species contact rates, and over time reduces the number of novel pathogens *P*_*i*_ to which each species is vulnerable (Eq. 5).

**Figure 2:**
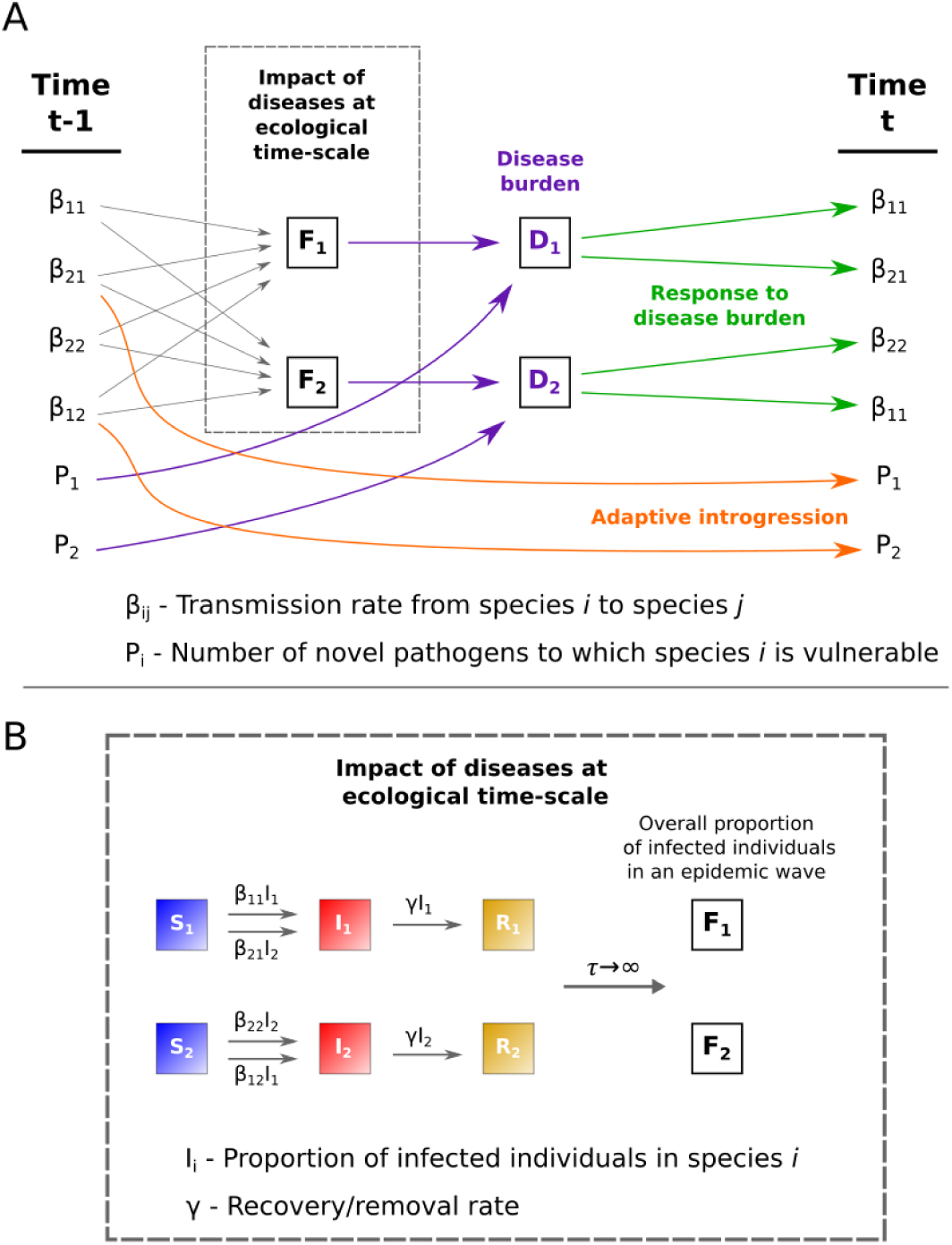
Schematic representation of the two-time-scales model, describing disease and introgression dynamics between two species. The figure describes the transition between two time steps at the evolutionary time scale, *t* − 1 and *t*. (A) Transmission rates (*β*_*ij*_, where for *i* = *j* these are within-species transmission, and for *i* ≠ *j* between-species transmission from species *i* to species *j*) determine the average impact of pathogens (*F*_*i*_) at the ecological time scale (dashed box, detailed in B), according to Eqs. 2 and 3. The pathogen package size (*P*_*i*_) and the average impact of each pathogen (*F*_*i*_) determine the disease burden (*D*_*i*_), in purple, according to Eq. 1. The species respond to disease burden by adjusting contact rates (*β*_*ij*_), in green, according to Eq. 4. Inter-species contact results in gene flow and adaptive introgression, reducing pathogen package sizes (*P*_*i*_), in orange, according to Eq. 5. (B) Impact at the ecological time scale (dashed box in A) is modeled in an SIR epidemic framework, as the average impact of an epidemic. Individuals transition from state S (susceptible) to state I (infectious) from either within-species infections or between-species infections (Eq. 2). Individuals in state I transition to state R (recovered/removed) at rate *γ* (Eq. 2). The impact of an epidemic (*F*_*i*_) is measured as the overall proportion of individuals in species *i* infected throughout the run of the epidemic (Eq. 3).

### 2.1 Symmetric conditions at the time of initial contact

We first characterize the general behavior of our model by exploring a scenario in which initial conditions are symmetric for the two species. An outcome of this scenario is presented in Figure 3. A more thorough exploration of the parameter space, with different parameterizations of the model, is presented in Supplementary Note A, with qualitatively similar results.

**Figure 3:**
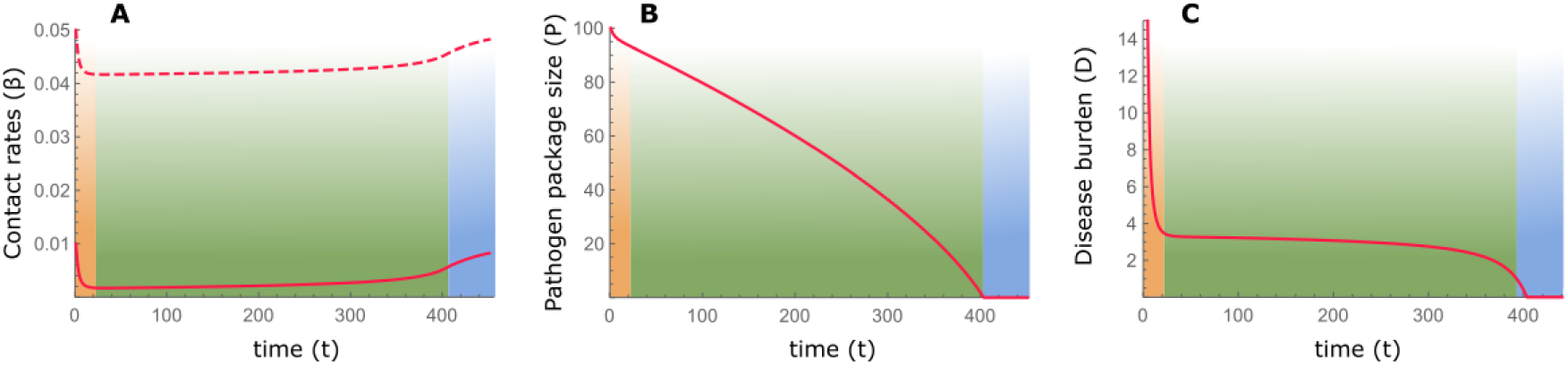
Disease-introgression dynamics between two species with symmetric initial conditions. The dynamics are described by the two-time-scales model (Fig. 2), following equations 1–5. The initial conditions are assumed to be symmetric for the two species (*β*_*ii*_ = *β*_*jj*_ and *β*_*ij*_ = *β*_*ji*_), and therefore, the dynamics for the two species are identical, indicated by red curves. (A) Between-species contact rates (*β*_*ij*_, *i* ≠ *j*) appear in solid curves, and within-species contact rates (*β*_*ii*_) appear in dashed curves (Eq. 4). (B) Pathogen package size (*P*_*i*_), measured in number or diversity of novel pathogens to which the species are vulnerable (Eq. 5). (C) Disease burden (*D*_*i*_), proportional to pathogen package size *P*_*i*_ and to the impact of diseases on each of the species, *F*_*i*_ (Eq. 1). Three phases are observed in the dynamics: (1) Initial response to heavy disease burden, with decreased contact rates (orange); (2) Long-lasting stable phase with low but steady levels of disease burden (green); (3) Destabilization following the release from disease burden and recovery to initial conditions (blue). The parameters for the scenario modeled are *β*_11_(0) = *β*_22_(0) = 0.05; *β*_12_(0) = *β*_21_(0) = 0.01; *γ* = 0.045; *P*_1_(0) = *P*_2_(0) = 100, with scaling parameters *a* = 5 *×* 10^−5^; *b* = 0.02; *c* = 100.

When the two species first come into contact, high contact rates and vulnerability to many novel pathogens result in high disease burden (Eq. 1), which elicits a rapid response of large effect, and the species lower both within- and between-species contact rates (Fig. 3, orange phase). Following this initial response, the species maintain stable but low contact rates for an extended period (Fig. 3A, green phase). During this period, the pathogen package *P*_*i*_ is reduced by adaptive introgression (Fig. 3B, green phase; Eq. 5), but disease burden is kept low and close to constant by the continuous minor adjustment of contact rates (*β*_*ij*_) made by the species (Fig. 3C, green phase; Eq. 4). We interpret the disease burden at this phase of the dynamics as limiting either species from expanding into regions dominated by the other, thereby leading to geographic localization of the interaction front. This localization is further reinforced by the tendency of bands to isolate themselves, as demonstrated by low contact rates (Fig. 3A, green phase).

Once disease burden is eventually removed due to introgression, the dynamics destabilize, and the species, released from disease burden, begin to recover and return to the initial state that existed before contact was made (Fig. 3, blue phase). This destabilization would then allow other dynamics, previously overshadowed by disease burden, to play out.

### 2.2 Asymmetric conditions at the time of initial contact

In reality, it is unlikely that conditions were symmetric at the time when Moderns and Neanderthals came into contact in the Levant (e.g., with respect to different pathogen co-evolution trajectories in Africa and in Eurasia). We therefore model asymmetric initial conditions, focusing for tractability on one possible aspect of asymmetry at a time. Figure 4 explores the effect of different initial pathogen package sizes (*P*_2_(0) > *P*_1_(0)), and asymmetry in initial contact rates is explored in Supplementary Note A.

**Figure 4:**
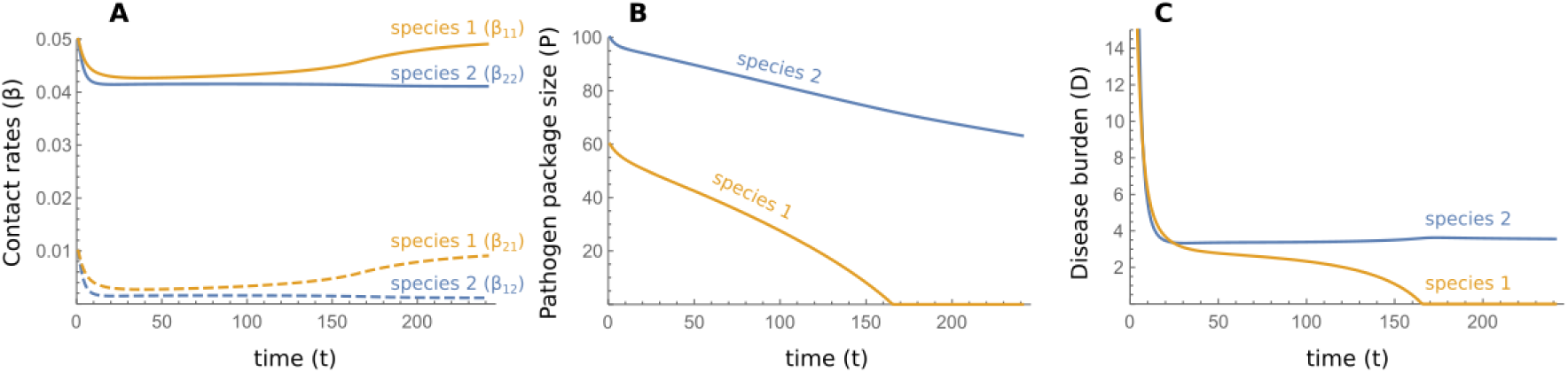
Disease-introgression dynamics between two species with asymmetric initial conditions. The dynamics are described by equations 1–5, with parameters identical to those in Fig. 3 except that species 1, in orange, is initially exposed to a smaller pathogen package than species 2, in blue, i.e. species 2 experiences more novel pathogens at the time of contact; *P*_1_(0) = 60, *P*_2_(0) = 100. (A) Between-species contact rates (*β*_*ij*_, *i* ≠ *j*) appear in dashed curves, and within-species contact rates (*β*_*ii*_) appear in continuous curves (Eq. 4). (B) Pathogen package size, *P*_*i*_. (C) Disease burden, *D*_*i*_. Species 1 experiences a lower disease burden at the time of contact, and therefore responds less strongly and maintains higher contact rates (in A). This higher contact rate leads to faster rates of adaptive introgression in species 1 (in B). Finally, species 1 is released from disease burden sooner than species 2 (in C), while still carrying many pathogens to which species 2 is vulnerable (in B).

When *P*_2_(0) > *P*_1_(0), species 2 experiences higher disease burden at the time of contact (*D*_2_(0) > *D*_1_(0)) due to the higher number of novel pathogens to which it is exposed (Eq. 1). This initial condition is reflected in a stronger initial response to disease burden, and lower within-species and incoming between-species contact rates (Fig. 4). The dynamics then enter a stable phase, similar to that in the symmetric case, but with some exceptions. First, species 1 retains slightly higher contact rates than species 2 during the stable phase (Fig. 4A), because the disease burden species 1 experiences is lower than that of species 2. Second, due to these higher contact rates, and specifically between-species incoming contact rates, the rate of adaptive introgression into species 1 is higher than into species 2 (*cβ*_21_(*t*) > *cβ*_12_(*t*)) during this period; Eq. 5). This pattern, in conjunction with the initial difference in pathogen package sizes, permits species 1 to overcome disease burden (i.e., reaching *P*_1_ = 0) earlier than species 2. Different parameterizations of the model yield similar qualitative results (Supplementary Note A).

These dynamics, which are qualitatively similar with other sources of asymmetries (Supplementary Note A), mean that the species that was initially less vulnerable, species 1, overcomes disease burden sooner, and is therefore released sooner from the disease limitation to enable expansion into regions dominated by the other species. At the time that species 1 is released from the novel disease burden, species 2 is still vulnerable to many novel pathogens carried by species 1 (Fig. 4B).

### 2.3 Feedback between disease transmission and introgression

Disease spread between species, in general, depends on the levels of within- and between-species contact rates, with higher contact rates resulting in greater disease impact. The level of contact between species, and specifically the amount of interbreeding, also determines the expected rate of gene flow, and higher contact rates are expected to result in more rapid adaptive introgression. That both of these phenomena — disease transmission and adaptive introgression — depend on inter-species contact rates, but have opposite effects on the contact rates, generates feedback that complicates prediction of disease and introgression dynamics (Fig. 2).

In our model, high disease burden for a species negatively impacts contact rates (−*a*_*ij*_*D*_*j*_ in Eq. 1), and lower between-species contact rates in turn negatively impact the rate of adaptive introgression for that species (−*cβ*_*ij*_ in Eq. 5). Therefore, this process generates a positive feedback that amplifies any initial differences in disease burden over time. In the case of asymmetric pathogen package sizes, the effect of differences in disease burden on the rate of adaptive introgression can be seen in the differing slopes of the pathogen package size trajectories of species 1 and 2 (Fig. 4B). Species 1, which at the time of contact experiences a lower disease burden, also adapts faster to the novel pathogens due to its maintenance of higher incoming between-species contact rates (*β*_21_ > *β*_12_; Fig.4A). In the scenario shown, this asymmetry results in the initial difference of 40 pathogens in the pathogen package sizes at the time of contact (the units could be different, such as pathogen diversity, but we focus on the relative outcomes without attention to the interpertation of the units), growing to about 75 pathogens by the time species 1 has overcome disease burden (Fig. 4B).

Another implication of this feedback is observed when species 1 is released from disease burden, i.e. reaches *P*_1_ = 0 (Fig. 4B). At this point, species 1 begins to increase its contact rates (Fig. 4A), and in so doing increases the impact of disease in the two-species system (Eq. 2 and 3). However, at this point only species 2 is vulnerable to the novel disease, and therefore only species 2 is forced to respond by further reducing contact rates (Fig. 4C), further slowing the rate of adaptive introgression into species 2. The consequence of the feedback is, therefore, that the species that overcomes disease burden sooner exerts an additional disease pressure on the other species during the destabilization phase.

### 2.4 Alternative model

The model presented above focuses on behavioral responses to disease burden. However, given that the model requires a seemingly complex behavior — a population must consciously or unconsciously modulate its interaction with other populations based on the amount of disease burden it experiences — it is worthwhile to consider if alternative models that emphasize other factors can produce similar dynamics. We explored an alternative model, which focuses on demographic processes and endemic diseases. In this model, disease dynamics, adaptive introgression, and population dynamics occur in parallel, over a single time scale (“single-time-scale model”), as described by equations 6–9. Here, we assume each species is entirely infected by a suite of endemic pathogens, to which it has evolved tolerance. The pathogens inflict no harm on the hosts in the source species, but they are novel to the other species, and therefore induce increased mortality (*α*_*i*_(*t*)) to infected individuals in the non-source species (*I*_*i*_(*t*)). In this model, population densities, *N*_*i*_(t), are determined by density-dependent growth rates and by mortality rates (Eq. 7). We assume that between- and within-species encounters are determined according to a process similar to Brownian-like motion of colliding particles, and therefore, contact intensities are modeled as being proportional to the population densities (Eq. 8). In this model, response to disease burden is determined purely through demographic effects. Adaptive introgression has the effect of decreasing the mortality rates associated with the novel pathogens, and the introgression rates, like the disease transmission rates, are proportional to the inter-species contact intensity (Eq. 9).

Under symmetric initial conditions, the dynamics are similar to those observed in the two-timescales model: an initial phase of high disease burden resulting in a decrease in population densities (Fig. 5A–C, orange phase), followed by a long phase of low but stable disease burden with low population densities (Fig. 5A–C, green phase), and eventual destabilization and population growth as disease burden is overcome (Fig. 5A–C, blue phase).

**Figure 5:**
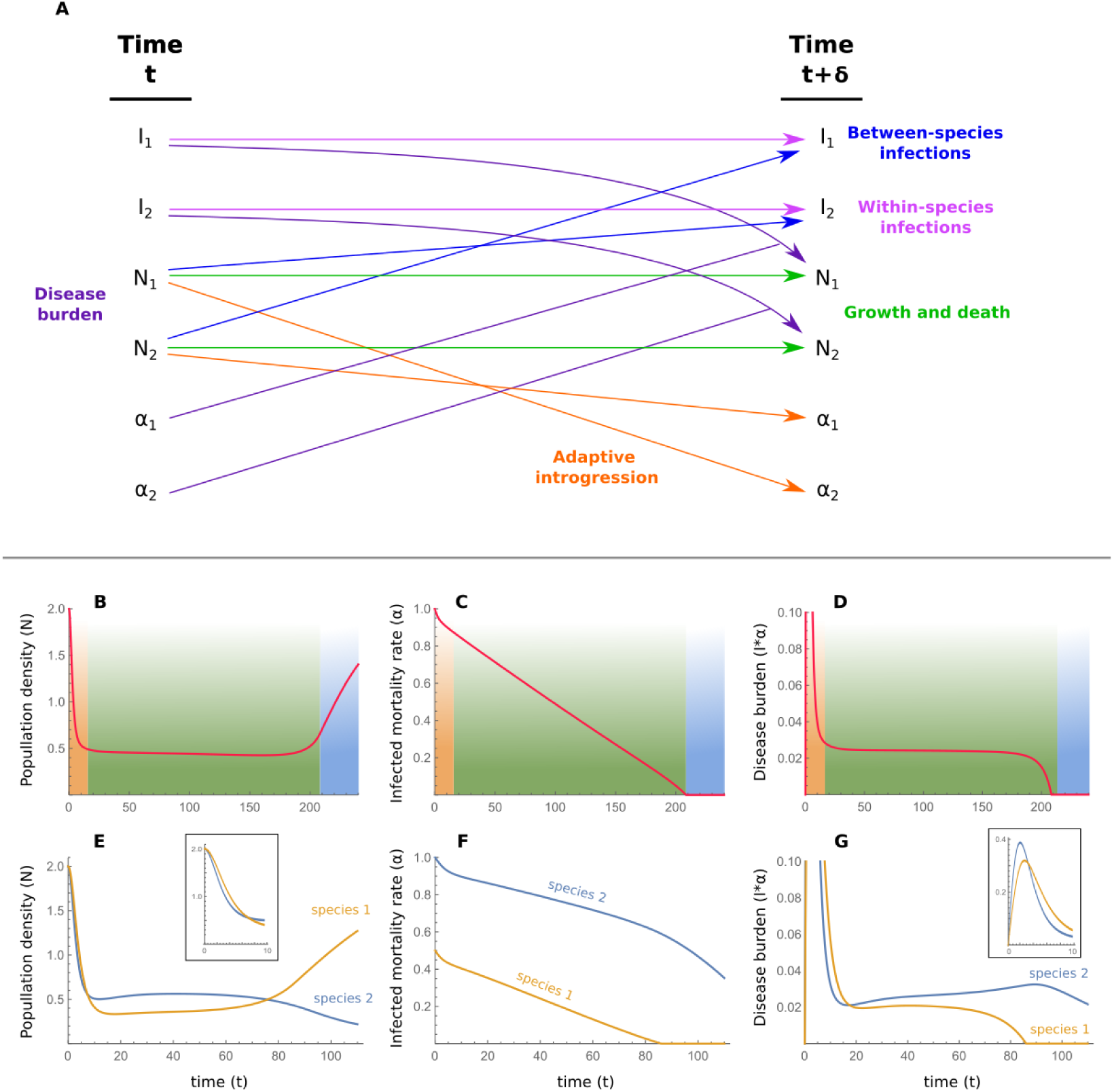
Alternative single-time-scale model of disease-introgression dynamics. (A) Schematic description of the model. The dynamics of model are described by equations 6–9. In this model, the population densities (*N*_*i*_(*t*)) are modeled, with disease burden caused by diseaseinduced mortality (*α*_*i*_(*t*)) of infected individuals (*I*). Adaptive introgression reduces disease-induced mortality over time. (B)–(D) show the dynamics with symmetric initial conditions, comparable to Figure 3, and (E)–(G) show results with asymmetric initial conditions, comparable to Figure 4. (B) and (E) show population densities, *N*_*i*_(*t*) (Eq. 7). (C) and (F) show the added mortality rate due infection from novel pathogens, *α*_*i*_(*t*) (Eq. 9). (D) and (G) show disease burden, measured as *I*_*i*_(*t*) *× α*_*i*_(*t*) (Eq. 8 and 9). The parameters for the modeled scenario are λ = 0.15; *K* = 1; *β*_11_ = *β*_22_ = 0.5, *β*_*a*_ = 0.1; *µ* = 0.05; *c* = 0.1. For the symmetric case, (B)–(D), *α*_1_(0) = *α*_2_(0) = 1, and for the asymmetric case, (E)–(G), species 1 experiences less disease-related mortality than species 2, *α*_1_(0) = 0.5 and *α*_2_(0) = 1. Inset graphs show the initial time periods in (E) and (G).

When the initial conditions are asymmetric with respect to the pathogen packages, *α*_2_(0) > *α*_1_(0), then species 2 suffers higher mortality due to novel pathogens than species 1. In this case, we observe that the two species initially respond strongly to disease burden and that population densities are reduced (Fig. 5D). A long stable phase of low population densities then follows. As is seen in the two-time-scales model, species 1 is released from disease burden sooner than species 2, when it begins to increase in density towards the initial density at the time of contact (Fig. 5D–F). With initial asymmetry in population densities (*N*_2_(0) > *N*_1_(0)), dynamics are qualitatively similar, with the initially less dense species 1 overcoming disease burden sooner in the scenarios examined (Supplementary Note A).

In the two-time-scales model, the separation of the ecological and evolutionary times-scales introduces some simplification to the dynamics, since the dynamics in the ecological time-scale are mediated to the evolutionary time-scale through a single summarizing parameter of disease impact (*F*_*i*_(*t*)). In the single-time-scale model, however, interaction between disease, introgression, and demography occurs on the same time-scale, resulting in more complex feedback between the different processes. Therefore, whether the consequences of the feedback are an increase or a decrease in asymmetries at the time that species 1 overcomes disease burden, depends on the source of the asymmetry and the parametrization of the model. In the scenario in Figure 5, and with many of the other scenarios we explored (Supplementary Note A), adaptive introgression in species 1 is more rapid than in species 2, and the differences between the disease-related mortality rates are increased at the time that species 1 overcomes disease burden, relative to the initial asymmetry; however, unlike in the previous model, this outcome is not always the case, and in some cases the differences in mortality rates between the species end up being lower at the time species 1 overcomes disease burden than at the beginning of the dynamics (e.g., Fig. S8K and Fig. S10K). Once species 1 is released from disease burden and increases in size, the feedback is similar to that in the twotime-scales model, since the increased population densities mean higher disease transmission in the two-species system, which at this point is detrimental only to species 2.

## 3 Discussion

Our results demonstrate that disease and introgression dynamics can explain the persistent stable phase of inter-species dynamics that preceded the replacement of Neanderthals by Moderns. They also suggest that such dynamics may have implications regarding the replacement phase.

### 3.1 Interpretation of the models

In the two-time-scales model, the adaptive response is in the contact rate itself, which is akin to assuming conscious or unconscious behavioral or cultural modification of contact rates [72]; for example, a band of hunter-gatherers may decrease inter-species contact by not accepting, or not taking by force, individuals of the other species. They may limit contact even further by avoiding other bands completely. However, reduction in contact rates in response to disease burden may also have occurred due to decreased population densities, and therefore this model may be viewed as one that incorporates implicitly the combined effect of both behavior and demography.

The model includes an implicit assumption regarding intra-species gene flow, which warrants elaboration. Immune-related alleles are assumed to spread rapidly once introgressed; that is, introgression relieving disease burden acts on the species as if they were cohesive units. Although this assumption may be inappropriate for large, geographically-structured, populations, our model is concerned only with the small peripheral region in which interaction occurs, and in which selection could have acted to favor beneficial immune-related genes. Under such conditions, the assumption that introgressed immune-related genes could spread rapidly in the contact zone appears reasonable.

We also assume that genetic adaptation via de novo mutations is negligible compared to adaptive introgression in reducing disease burden. This assumption amounts to an assumption that introgression of pre-adapted alleles through interbreeding occurs more frequently than the appearance of novel mutations conveying resistance to a disease, under the scenario of Neanderthal-Modern interaction. We explore incorporation of regular adaptation into the models in Supplementary Information Note B, with results qualitatively similar to those presented in Figures 3–5 but with shorter stable phases.

In our alternative single-time-scale model, the adaptive response is a reduction in the level of disease-induced mortality, which is akin to assuming evolution of tolerance to the invading pathogen package. These exclusively demographic, rather than behavioral, dynamics can be readily interpreted in the context of an immunological or physiological response to assaults from novel pathogens. The model also assumes rapid within-species assimilation of acquired tolerance, similar to the assumption in the two-time-scales model. Modeling of regular adaptation in the single-time-scale model is addressed in Supplementary Note B. Competition for resources between the species in the contact-zone may have demographic consequences, but is not incorporated into the model; however, we have added competition to our model in Supplementary Note C, and our analysis again yields similar results to those in Figure 5 but with longer stable phases.

We have shown here that disease dynamics are sufficient to explain the long period in which the contact-zone was confined to the Levant; however, other factors may have played a role. One such factor may be the species adaptation to their respective environments at the core of their geographical ranges [9, 11, 73]. Many such adaptations have been proposed, ranging from morphology that supports faster or slower heat loss [55, 74, 75] to physiological traits that support different hunting methods [57]. Such adaptations could have limited migration of bands of one species into regions of the other species, similar to the effect of disease burden. However, local adaptation by itself would not explain the rapid destabilization of the interaction front, since the period of 50-40kya was not characterized by unprecedented environmental change that in itself would have provided Moderns an advantage across the entirety of the Neanderthals range [76–79] but see [80–82] and the discussion in [83]. Similarly, no clear evidence has been found so far of a cultural shift among Moderns that would have provided them a sudden advantage over the Neanderthals [84].

### 3.2 Implications for the replacement of Neanderthals by Moderns

When the initial conditions are asymmetric, one species apears to acquire tolerance to novel diseases sooner than the other (Fig. 4 and 5D–F), permitting that species to expand its range earlier. A particularly plausible source of asymmetry, in the case of Moderns and Neanderthals, is the differences in pathogen complexes to which each of the species was adapted. Biotic diversity, on many taxonomic scales, is higher in the tropics [85, 86], including in human pathogens [61, 62]. In the Levant, where climate was intermediate between the temperate and tropical zones [87], many pathogens carried by both species would have been potential sources for diseases. It is therefore possible that Neanderthals would have had to adapt to a larger number of pathogens than did Moderns (*P*_2_(0) > *P*_1_(0) or *α*_2_(0) > *α*_1_(0) in our models, with species 1 representing Moderns and species 2 representing Neanderthals), leading to earlier Modern release from disease burden (Fig. 4 and Fig. 5D–F).

Although we have not modeled spatial structure in the two species, following destabilization, bands of Moderns moving deeper into Neanderthal regions would have encountered bands of Neanderthals whose lineages had not interacted with Moderns, and who had been far enough from the long-standing front of interaction to not have received immune-related adaptive introgressions. These bands would thus have been even more vulnerable than Neanderthals in the Levant to pathogens carried by Moderns. The scenario is analogous to more recent events, such as when Europeans arrived in the Americas in the 15th and 16th centuries with a more potent pathogen package than that of the local inhabitants, not because of climate but because of higher population densities and contact with domesticated animals [88]. The colonization of the Americas was followed by rapid replacement of Native Americans, facilitated by disease spread [63–71, 88].

That diseases played an important role in the inter-species dynamics of Moderns and Neanderthals [53, 59, 60] is suggested by studies that compare Neanderthal genomes with current-day Modern genomes, and that argue that genomic regions relating to disease immunity and tolerance are enriched in introgression of Neanderthal genes [2, 43, 47, 49–53]. This result suggests that introgression was adaptive, and that diseases were a significant enough burden that natural selection in Moderns favored introgressed lineages that included immune-related genes. Interestingly, recent indications from analysis of European Neanderthal genomes suggests that gene flow was not symmetric between the species, and that more genes were introgressed from Neanderthals into Moderns than in the other direction [89]. This asymmetry is predicted by our two-time-scale model, which addresses differential inter-species contact intensities (Fig. 4A, dashed curves); note, however, that the Neanderthals analyzed in producing this evidence of assymytry were not sampled in the Levant [89].

### 3.3 Future investigation of the Neanderthal-Modern disease landscape

Further investigation of the role of diseases in the interaction between Neanderthals and Moderns would require a better understanding of the pathogen landscape during this period. One direction is the study of the genomic regions that were under selection at the time of contact [2, 43, 47, 49– 53]. A few recent phylogenetic studies of pathogen have indicated that some pathogens might have been transmitted from Neanderthals or other archaic humans to Moderns [45–48]. A particularly useful direction for elucidating the disease landscape would be genomic studies [44, 90–96] of ancient pathogens recovered from archaeological Neanderthal and Modern sites.

The consequences of inter-species disease dynamics may be evident in archaeological findings as well. For example, as suggested by our model, disease burden in the Levant might have affected population densities (Fig. 2D and Fig. 4D), reducing them compared to those in adjacent regions where novel pathogens have not yet been introduced, to those prior to contact, or to those after release from the disease burden. Population densities might be estimated by the assessment of archaeological site density and site complexity [97–99]. Population density can potentially also be assessed via analysis of resource exploitation; for example, mean prey size may reflect predation pressure, allowing estimation of hominin population density [100–102]. Further archaeological studies that target parameters of population demography in the Levant at the time in question will be important to test the predictions of our models.

### 3.4 Conclusion

A major focus in the study of the inter-species dynamics between Moderns and Neanderthals has been the relative rapidity with which Moderns replaced Neanderthals across the majority of Eurasia. In this study, we suggest that analyzing the phase that preceded the eventual replacement is valuable as well. That the two species’ front of interaction was constrained to the Levant for tens of thousands of years is puzzling, particularly in light of the short time — a few millennia — within which the replacement across the rest of Eurasia was completed. We have drawn insights from the field of disease ecology to suggest that disease dynamics may explain the long period of stability that preceded the replacement. We have explored this possibility using mathematical modeling, deriving predictions can inform future exploration. We propose that this approach provides insights into the inter-species dynamics at the transition between the Middle and Upper Paleolithic periods, particularly due to the sparsity of the material record from this period and in consideration of the promise that DNA sequencing and dating technologies hold. Such modeling, coupled with new technologies and with novel approaches in prehistoric archaeology, may act synergistically to allow a new interpretation of this exciting period in human evolution.

## Supporting information

## 4 Acknowledgments

We would like to thank David Gokhman, David Freisem, David Enard and Anna Belfer-Cohen for insightful comments. GG and OK were funded by fellowships from the Stanford Center for Computational, Evolutionary and Human Genomics (CEHG). OK was also funded by a grant from the John Templeton Foundation. This project was supported by NSF grant BCS-1515127 awarded to NAR.

## 5 Author contributions

GG and OK conceived the study. GG, WMG and OK developed the models. NAR, MWF and EH contributed ideas to the conceptual framing of the project. GG, OK and WMG drafted the paper. All authors read and commented on the manuscript.

## 6 Methods

We develop a modeling approach to explore the effects of disease and introgression in a two-species system. In the two-time-scales model, the evolutionary and ecological times scales are separated (see [103–107] for other such models). On the ecological time scale, the spread of diseases is modeled, and on the evolutionary time scale, the species’ response to disease burden and the effects of introgression are modeled (Fig. 2). We assume that each species initially carries a pathogen package with which it had co-evolved, and to which it is therefore tolerant. A species suffers no significant negative impact from its associated pathogen package, but its pathogens are novel to the other species, which is vulnerable to them and is harmed considerably.

In the single-time-scale model, a similar scenario is modeled, and a single time scale is used to model ecology, disease, and introgression as interacting parallel processes. The process that drives the response to disease burden is a demographic process.

### 6.1 Two-time-scales model

On the evolutionary time scale, we model disease burden, response to disease burden, and introgression using a discrete-time model (Fig. 2). Each time step in this model (*t*) is assumed to be large enough for disease processes to be experienced by the populations and for the populations to respond to these disease pressures. For simplicity, we focus on disease dynamics described by within- and between-species contact rates (*β*_*ij*_ for contact rates in which pathogens from species *i* are transmitted to species *j*). We model response to disease burden implicitly as adjustments of these rates.

Initially, each species *i* experiences a novel pathogen package of size *P*_*i*_(0). This package size can be interpreted as the number, or diversity, of novel pathogens in species *j* (*j* ≠ *i*) to which species *i* is still vulnerable, the genes providing immunity to these pathogens not having yet introgressed into species *i*. At each time point, the disease burden experienced by the pair of species is modeled as:

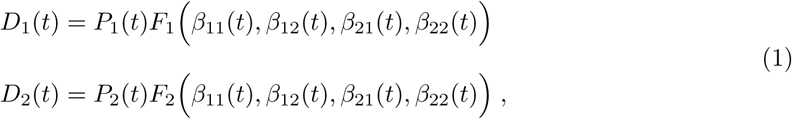

where *D*_*i*_(*t*) is the disease burden experienced by species *i* at time *t*, and *F*_*i*_(*t*) is a disease-ecology model describing the average impact of a single pathogen on population *i* given the contact rates *β*_*ij*_(*t*). *F*_*i*_ describes the faster ecological time scale in the model.

We model the ecological process using a two-species well-mixed SIR epidemic model [108–111] (Fig. 2B). In an SIR model, individuals can be in one of three states — susceptible (*S*), infectious (*I*), and recovered/removed (*R*), which are measured in terms of their proportion in the entire population (i.e. *S* + *I* + *R* = 1). In addition to transmission rates (*β*_*ij*_), the SIR model requires additional parameters describing the recovery/removal rates for the two species, *γ*_1_ and *γ*_2_ for species 1 and 2, respectively; these parameters represent rates of either recovery with immunity or death from the disease, which have similar outcomes in their effect on spread of the disease, because in both cases the host can no longer transmit the disease. For simplicity, we assume symmetry between the species in recovery/removal rates, i.e. *γ* = *γ*_1_ = *γ*_2_. The equations governing the dynamics of the SIR model over time, which we term *τ* to distinguish from the longer evolutionary time scale *t*, are:

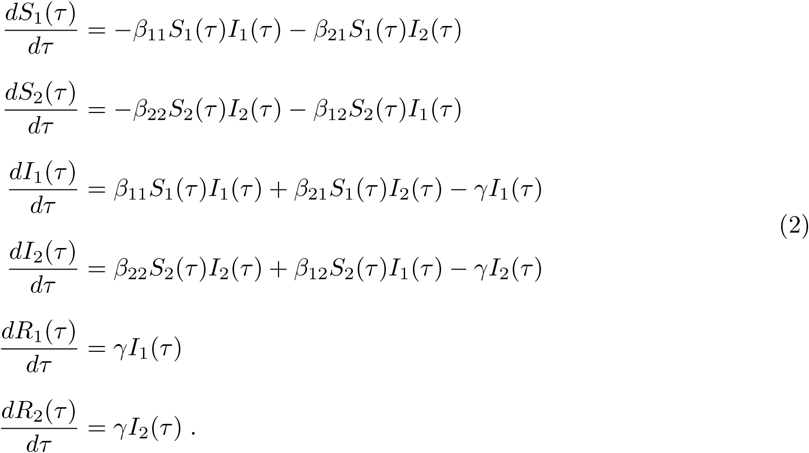

Note that at this time scale the *β*_*ij*_ parameters are considered fixed, and they change only at the evolutionary time scale. Individuals of species *i* in the susceptible state, *S*_*i*_, can transition to the infected state *I*_*i*_ by being infected either by individuals from their own species, at rate proportional both to *β*_*ii*_ and to the proportion of contacts between susceptible and infected individuals, or, similarly, by being infected by individuals from the other species at a rate proportional to both *β*_*ji*_ and the proportion of inter-species contacts between infected and uninfected individuals. Infected individuals transition to the recovered/removed state at a fixed rate, *γ*.

Under this SIR model with specific values of *β*_*ij*_ and *γ*, the impact of an average epidemic wave on a species, *F*_*i*_ for species *i*, is measured by the proportion of the population that was infected during an entire run of the epidemic:

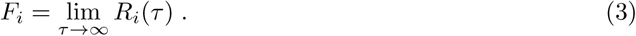

We measure this proportion in the case that the epidemic originates in relatively few individuals in species *j* (*I*_*i*_(0) = 0 and *I*_*j*_(0) = 0.01). The limit is taken over *R* since infected individuals inevitably end up in the *R* state, and therefore, tracking the eventual number of recovered/removed individuals amounts to tracking the number of overall infected individuals. The scenario modeled using an SIR model at the ecological level is one in which each species experiences repeated independent waves of epidemics originating in the other species. The epidemic spreads in the combined system of the two species, but only the non-source species is vulnerable to the effects of the disease. Consequently, the spread of disease, in our model, has no impact on the source species, except in its effect on disease spread in the non-source species.

On the evolutionary time scale, after experiencing disease burden (e.g. repeated epidemic outbreaks that originated in the other species), the species modify their contact rates accordingly. Such modifications can be behavioral in nature, either via a conscious process, whereby individuals notice that actively reducing within-species and particularly between-species contact rates reduces the impact of outbreaks, or via unconscious alteration of behavior, involving long-term selection on cultural traits, or instinctive factors such as stress-induced aversion to strangers. Contact rate modification can also be demographic in nature, where disease burden reduces population densities and increases the geographic distances between groups of individuals.

Each species is also assumed to have initial intrinsic within- and between-species contact rates, which reflect the typical contact rates unhindered by the novel pathogens. We model the adjustment of contact rates as being proportional to the disease burden, countered by the tendency of species to return back to their intrinsic behavioral and demographic states. These tendencies are modeled as proportional to the difference between the current state and the original state. Therefore, adjustment of contact rates in response to disease burden is modeled as follows:

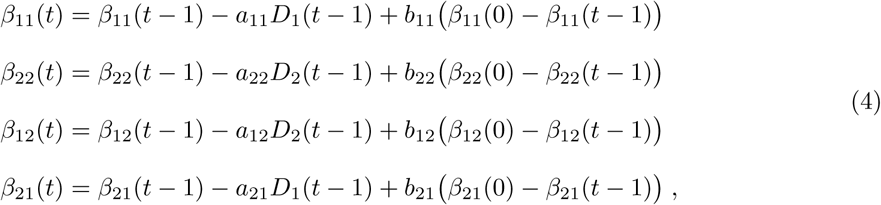

where the *a*_*ij*_ are parameters scaling the response to disease burden and the *b*_*ij*_ are parameters scaling the tendency to return to the initial contact rates. For simplicity, we assume symmetry for these parameters, and define *a* = *a*_*ij*_ and *b* = *b*_*ij*_ for all *i* and *j*.

Finally, we model the effect of introgression on the number of novel pathogens experienced by the species, assuming that it is proportional to inter-species disease transmission:

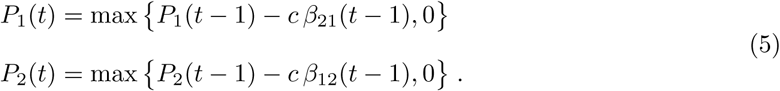

Here, *c* is a parameter scaling the rate of adaptive introgression relative to the rate of disease transmission. Since these two rates are defined at different times scales (disease transmission on the ecological time scale in Eq.4, and introgression on the evolutionary time scale in Eq.5), *c* has an additional role in the models as the parameter that scales the two times scales.

In equation 4, the response to disease burden is modeled as a reduction in within-species and incoming between-species rates. This choice is particularly appropriate for behavioral responses to disease pressure, where between-species contacts are most likely determined by the willingness of the receiving population to accept contact.

### 6.2 Single-time-scale model

In this section, we describe a continuous-time model of an endemic disease process that occurs in parallel to immune-related introgression and demographic dynamics. The model is a birth-death SI model [110] in which population sizes are affected by the birth and death rates, and disease burden is modeled as an increased death rate in individuals infected by novel pathogens (*α*_*i*_(*t*)) over the natural death rate (*µ*). The model tracks the species population densities (*N*_*i*_(*t*)) given intrinsic growth rates (λ). In order for growth rates to be density-dependent, the actual growth rates are given as a function of *N*_*i*_ [112, 113]:

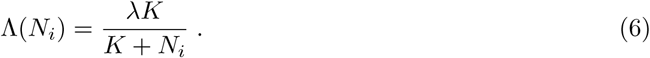

Here, λ is the maximal growth rate (achieved in sparse populations), and *K* is a half-maximum growth rate density parameter. In other words, 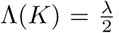; populations of density *K* grow at half the maximum growth rate. For simplicity, we assume that the intrinsic growth rate (λ), the half-maximum growth rate density parameters (*K*), and the natural mortality rates (*µ*), are the same for both species.

Population dynamics therefore reflect population growth, natural population mortality, and the additional mortality incurred due to the novel pathogens, where the number of individuals infected by novel pathogens is *I*_*i*_(*t*):

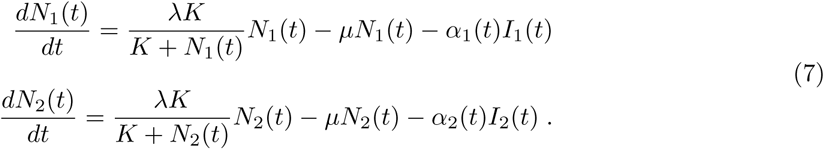

Initially, prior to inter-species contact, we assume no individuals are infected by novel pathogens (*I*_*i*_(0) = 0), and that the populations are at demographic equilibrium, implying that 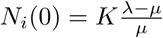 (see Supplementary Note A for different assumptions).

Next, we model disease dynamics in the populations using an SI model, in which each individual can only move from a susceptible state to an infected state, and cannot recover. The susceptible individuals (*N*_*i*_(*t*) − *I*_*i*_(*t*) in this model) can either be infected by any other individual from the other species, or by infected individuals in its own species. Note that we assume that all individuals in the source population are infected; this assumption is plausible for an endemic pathogen that has co-evolved with the species, and for which that species has little physiological response. Under the SI model, infected individuals can be removed only when they die, at rate *µ* + *α*_*i*_(*t*). The dynamics of the disease are therefore modeled by:

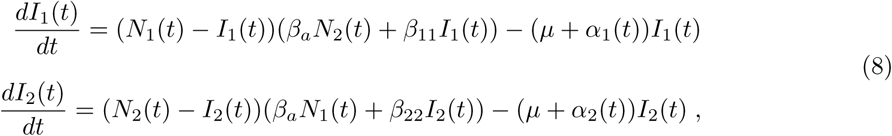

where *β*_11_ and *β*_22_ are the within-species transmission parameters and *β*_*a*_ is the symmetric betweenspecies transmission parameter, which are all proportional to the so-called “contact rate” (the rate at which individuals are close enough together to facilitate a pathogen transmission event) and the probability of transmission per contact [114]. Note that here, inter-species transmission is modeled as being driven purely by random encounters, affected only by the population densities. Hence, there is a single parameter for inter-species transmission (*β*_*a*_), unlike in the two-time-scales models, which has one for each direction of transmission (*β*_*ij*_ ≠ *β*_*ji*_ for *i* ≠ *j*). Additionally, contact rates in this model are fixed (i.e. they do not change throughout the dynamics) and are not subject to behavioral alteration in response to disease burden.

Finally, we describe the effect of transmission between the species on introgression. Here, disease burden is modeled not as a function of the number of pathogens, but more implicitly as the cumulative effect expressed by increased mortality rate (*α*_*i*_). Therefore, we model immune-related introgression as reducing the mortality associated with being infected by the novel pathogens, factored by the transmission intensity (force of transmission) between the species

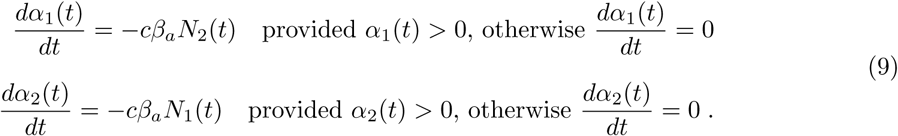

*c* is a scaling parameter interpreted as the ratio between immune-related gene transmission and disease transmission.

Equations 7–9 describe a simple SI model of endemic diseases in two species, where response to disease burden is purely demographic, and in which contact rates between the species are not affected by disease burden.

### 6.3 Numerical analysis

All systems of partial differential equations were solved numerically using the Mathematica software [115]. These numerical solutions were derived deterministically from explicitly stated initial conditions.

