## Supplementary material for "Disease and introgression explain the long-lasting contact zone of Modern Humans and Neanderthals and its eventual destabilization"

### Supplementary Notes

- A — Exploration of parametrization
- B — Modeling regular adaption in addition to adaptive introgression
- C — Modeling inter-species competition

### A Exploration of parametrization

In this section, we explore several of the parameters of the models. The goal of this section is not an exhaustive exploration of the parameter space, but rather to demonstrate that the dynamics are qualitatively similar for a significant portion of this space, in the sense that the three phases described in the main text (Figs. 3–5) can be observed. It is also aimed at providing some intuition about how the parameters control the dynamics, which we summarized as the time of onset of the second and third phases. We do not conduct this analysis for all the parameters in the models, but rather focus on a few which we believe such an analysis can provide insights.

For the two-time-scales model (Eqs. 1–5), we explore varying initial (which is also intrinsic) between-species contact rates ($\beta_{12}(0)$ and $\beta_{21}(0)$), the ratio between disease transmission and introgression ($c$), and initial asymmetry in within-species contact rates ($\beta_{11}(0) \neq \beta_{22}(0)$). We do not conduct the analysis for the scaling parameters ($a_{ij}$ and $b_{ij}$) which adjust the relative impacts of disease burden and tendency to return to initial conditions, since we vary the other parameters which determine disease burden. We also do not conduct the analysis for the recovery rate $\gamma$, which affects disease burden in a fairly straightforward way (longer recovery times amount to more infected individuals and higher $F_i$ in Eq. 3).

For the single-time-scale model, we explored varying between-species contact rates ($\beta_a$), the ratio between disease transmission and introgression ($c$), and initial asymmetry in intrinsic population densities ($N_1(0) \neq N_2(0)$). Since we focus on the disease and introgression processes, we do not vary the demographic parameters $\lambda$, $K$ and $\mu$.

For tractability, we vary one parameter at time, using the parameters presented in the main text as a baseline.

#### A.1 Two-time-scales model

In order to compare the dynamics over several parameter ranges, we summarized the dynamics using two values, the time of onset of phase II, and the time of onset of phase III (Figs. 3 and 5). For the time of onset of phase II, we first defined for each of the contact rate parameters the time point, $u_{\beta_{ij}}$, as the first time-step in which the absolute difference between $\beta_{ij}$ at consecutive time steps was lower than $10^{-3}$; in other words, $u_{\beta_{ij}}$ was defined as the lowest $t$ for which $|\beta_{ij}(t) - \beta_{ij}(t-1)| < 10^{-3}$. This is a discrete equivalent for finding the first time point in which the first derivative of the contact rate is very close to 0, indicating arrival at the stable phase. The threshold ($10^{-3}$) was selected so as not to be too small and miss the stabilization of the dynamics at phase II, but not too high as to define stability when the slope was still observably significantly different then 0. The overall onset of phase II, $u$, was taken to be the maximum over the four $u_{\beta_{ij}}$ values (for $i, j = 1, 2$), meaning that we consider the onset of phase II as the first time step in which all contact rates are considered to have arrived at the steady state.

The time of onset of phase III, $w$, was defined as the first time step for which disease burden in either species was removed; in other words, the earliest $t$ for which either $D_1(t) = 0$ or $D_2(t) = 0$.

##### A.1.1 Between-species contact rates

We first varied the between-species contact rates, which directly affects both the transmission of disease in the system and the rate of adaptive introgression. We used the parameter values in Figures 3 and 4 as a baseline, and varied $\beta_{ij}$ $(i \neq j)$ from 0.005 to 0.03 in increments of 0.001 (maintaining $\beta_{ij} = \beta_{ji}$). For between-species contact rates below 0.005, the numerical solutions of the differential equations in Eq. 2 were affected by computational precision errors due to very low values, and we therefore restricted our range to avoid these low ranges of transmission. We conducted this analysis for the symmetric $(P_1(0) = P_2(0))$ and asymmetric cases $(P_1(0) \neq P_2(0))$, as presented in the main text.

Figure S1 shows the time of onset for the two phases for the different within-species contact rates. We also show the full dynamics over time for two examples, one with between-species contact rates higher than those shown in the main text ($\beta_{ij} = 0.2$, in blue) and one with lower rates ($\beta_{ij} = 0.05$, in green). This we do to demonstrate that the dynamics are qualitatively similar to the scenarios described in Figures 3 and 4, in the sense that three distinct phases can be observed (Fig. S2).

The onset of phase II increases with increased between-species contact, for both the symmetric and asymmetric scenarios examined (Fig. S1A–B), while the onset of phase III decreases for both scenarios (Fig. S1C–D). Overall, since the changes in $w$ are much larger than those in $u$, the length of Phase II $(w - u)$ is more affected by $w$, and therefore, for the parameter range we examined, the length of phase II decreases with increased between-species contact. We also show two examples of the dynamics (Fig. S2), which show qualitatively similar dynamics to those in Figures 3 and 4, but the dynamics occur over different times scales, with faster dynamics in the case of high between-species transmission (in blue) compared to that with low between-species contact rates (in green).

The between-species contact rate parameter $\beta_{ij}$ plays two roles in the model: (1) Increase in $\beta_{ij}$ is expected to increase $F_i$ (Eqs. 1 and 2), and therefore increase disease burden, $D_i$ (Eq. 3), which would lead to a decrease in all contact rates (Eq. 4);(2) Increase in $\beta_{ij}$ increases the rate of introgression (Eq.5). At the early stages of the dynamics, when the species are strongly impacted by epidemics, increased disease burden results in significant reductions in between-species contact,and therefore slower introgression, leading to later onset of phase II (Fig. S1A and S1B). However, once disease burden decreases over time, the high intrinsic $\beta_{ij}(0)$ leads to faster introgression, reducing the time to onset of phase III (Fig. S1C and S1D).

##### A.1.2 Ratio of introgression to disease transmission

The two-time-scales model assumes that both introgression and disease transmission are coupled with the between-species contact rates, with a fixed parameter describing the ratio between introgression and between-species contact, $c$. We varied $c$ between 10 and 5000 in increments of 10, with the other parameters kept the same as those in Figures 3 and 4.

Fig. S3 shows the time of onset for the two phases for the $c$ values. We also show the full dynamics over time for two examples (Fig. S4), one with $c$ value higher than the one described in the main text ($c = 1000$, in blue), and one with a lower $c$ value ($c = 10$, in green). These are shown to demonstrate that the dynamics are qualitatively similar to the scenarios described in Figures 3 and 4, in the sense that three distinct phases can be observed.

The onset of phase II is unchanged for up to about $c = 1000$, above which the onset time $u$ decreases as the ratio $c$ increases (Fig. S3). The onset of phase III decreases with high $c$ values (Fig. S3), which is due to the fact that a high $c$ value means faster adaptive introgression and faster overcoming of disease burden, in both the symmetric and asymmetric cases.

The ratio between disease transmission and introgression, $c$, plays a role in Eq. 4, where higher $c$ values lead to faster introgression. Therefore, increasing $c$ results both in earlier onset of phase II and earlier onset of phase III (Fig. S3).

##### A.1.3 Initial asymmetry in contact rates

In Figure 4 in the main text, we explore the consequences of initial asymmetry in pathogen package size, $P$. Here we consider initial asymmetry in the intrinsic within-species contact rate, $\beta_{11}(0)$ and $\beta_{22}(0)$. Such asymmetry could reflect, for example, differences in population densities (within-species contacts occur at lower rates in sparser populations), but also differences in contact tendencies of the two species. One scenario where such differences may have occurred is if the incoming Moderns were characterized, over a long period, with lower population densities than the resident Neanderthals. However, this phase may have been a short transient phase for the Moderns, in which case it would not necessarily have been influential for the dynamics we consider.

To explore this source of asymmetry, we considered the symmetric case in Fig. 3 as a baseline and varied the within-species contact rates for species 2, $\beta_{22}(0)$, from 0.005 to 0.05 at increments of 0.001, with the other parameters kept the same as those in Figure 4 (in particular, $\beta_{11} = 0.05$). Figure S5 shows the time of onset for the two phases for the $\beta_{22}(0)$ values. We also show the full dynamics over time for two examples (Fig. S6), one with $\beta_{22}(0) = 0.03$ (in blue), and one with $\beta_{22}(0) = 0.015$ (in green).

With asymmetry in within-species contact rates, the time of onset of phase II is almost unchanged for the parameter range we investigated, but the onset of phase II is slightly sooner with more accentuated asymmetry (Fig. S5). Interestingly, and perhaps counterintuitively, with such asymmetry, species 2, which initially had lower within-species contact rates, overcomes disease burden sooner than species 1. This is because the lower within-species contact rates translate to lower $F_2$ values in Eq. 1. and hence lower initial disease burden (Fig. S6C and S6F), which allows it to maintain higher incoming between-species contact rates (Eq. 4, and Fig. S6A and S6D). These between-species contact rates are then translated to higher rates of adaptive introgression (Eq. 5, and Fig. S6B and S6E), and more rapid overcoming of disease burden.

The initial condition of $\beta_{22}$ determines the intrinsic within-species contact rates in the species (Eq. 4). In the model, within-species contact rates play a role in determining the impact of diseases on the species (Eqs. 1–3), but do not directly affect introgression. Therefore, decreasing the within-species contact rate of one species relative to the other, results in less impact of disease, allowing the species to maintain higher between-species contact rates and increases the rate of introgression. This leads to earlier onset of phase III with lower within-species contact rate in one of the species.

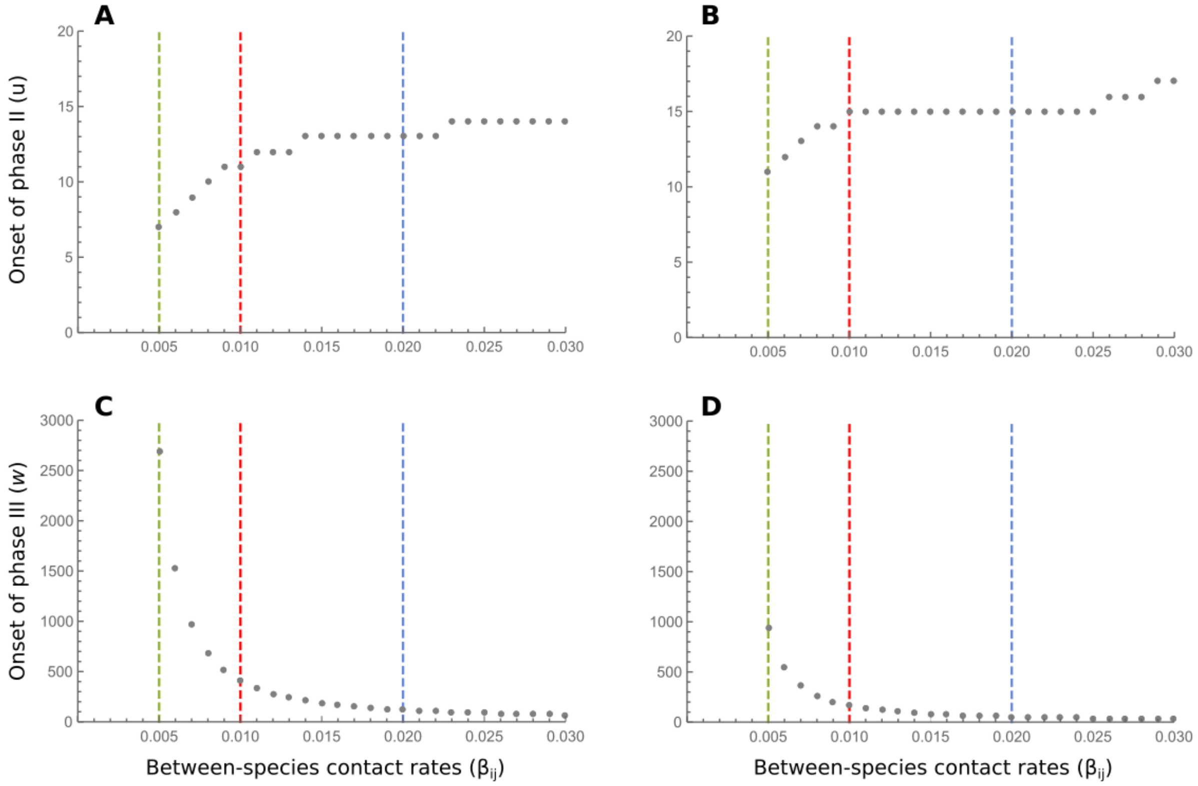


**Figure S1: Time of onset of phases II ($u$) and III ($w$) for different between-species contact rates ($\beta_{ij}(0)$) in the two-time-scales model.** Dashed lines show the parameter values for which we show the dynamics explicitly, in the main text in Figs. 3 and 4 (red), or in Fig. S2 (green and blue). (A) Onset of phase II for the symmetric case. (B) Onset of phase II for the asymmetric case ($P_1(0) = 60$, $P_2(0) = 100$). (C) Onset of phase III for the symmetric case. (D) Onset of phase III for the asymmetric case ($P_1(0) = 60$, $P_2(0) = 100$). Other than the the between-species contact rates, all other parameters are the same as those in Fig. 3 (for (A) and (C)) and Fig. 4 (for (B) and (D)).

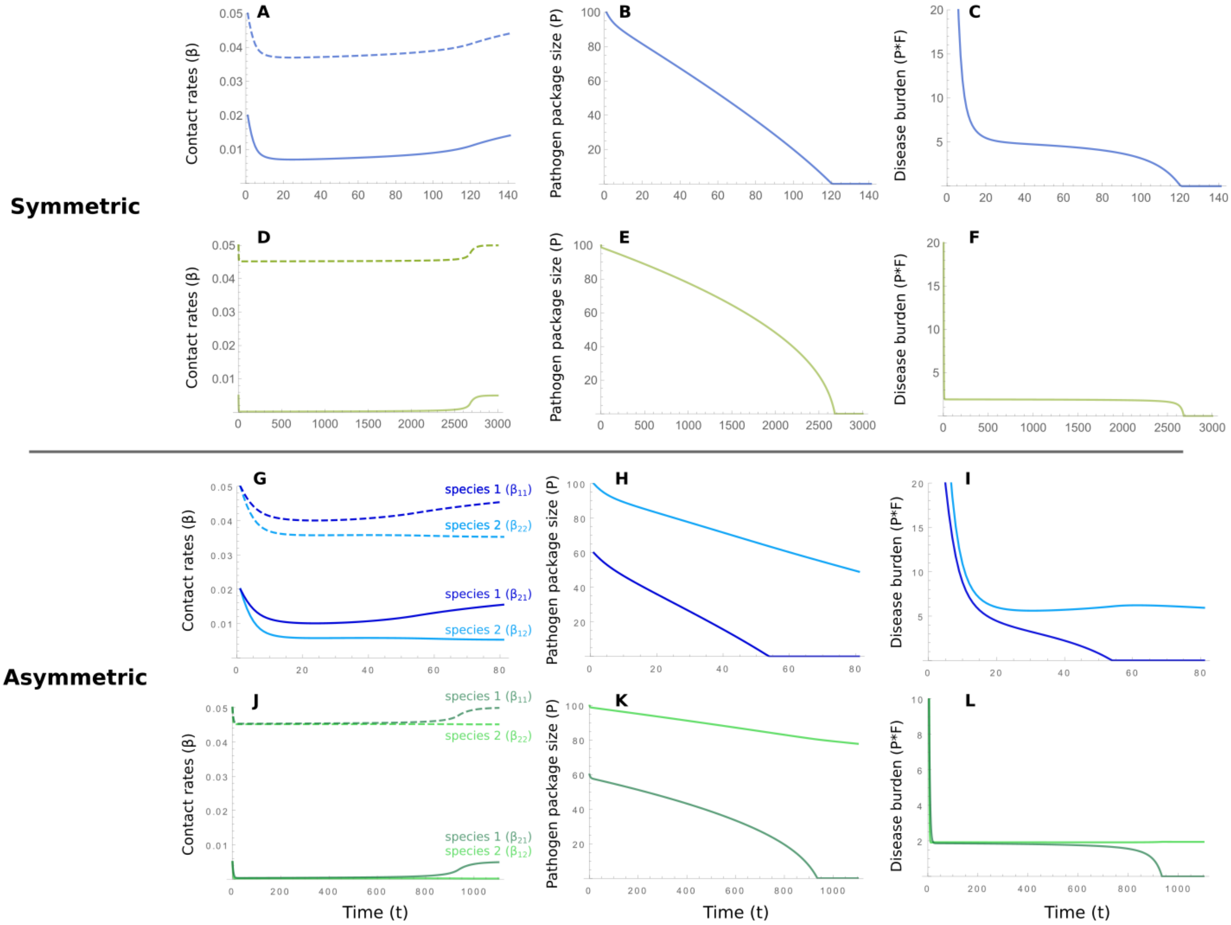


**Figure S2: Two examples of dynamics of the two-time-scales model for the values of between-species contact rates ($\beta_{ij}$) marked in Fig. S1**. (A)–(F) show the results for the symmetric case, comparable with Fig. 3, and (G)–(L) show the results for the asymmetric case, comparable with Fig. 4. For the panels with blue curves (corresponding to the blue line in Fig. S1), between-species contact rates are $\beta_{ij}(0) = 0.02$, while the rest of the parameters are the same as those in Fig. 3 and Fig. 4. For the panels with green curves (corresponding to the green line in Fig. S1), between-species contact rates are $\beta_{ij}(0) = 0.005$, while the rest of the parameters are the same as those in Fig. 3 and Fig. 4. Notice that the times scales for the different panels are different.

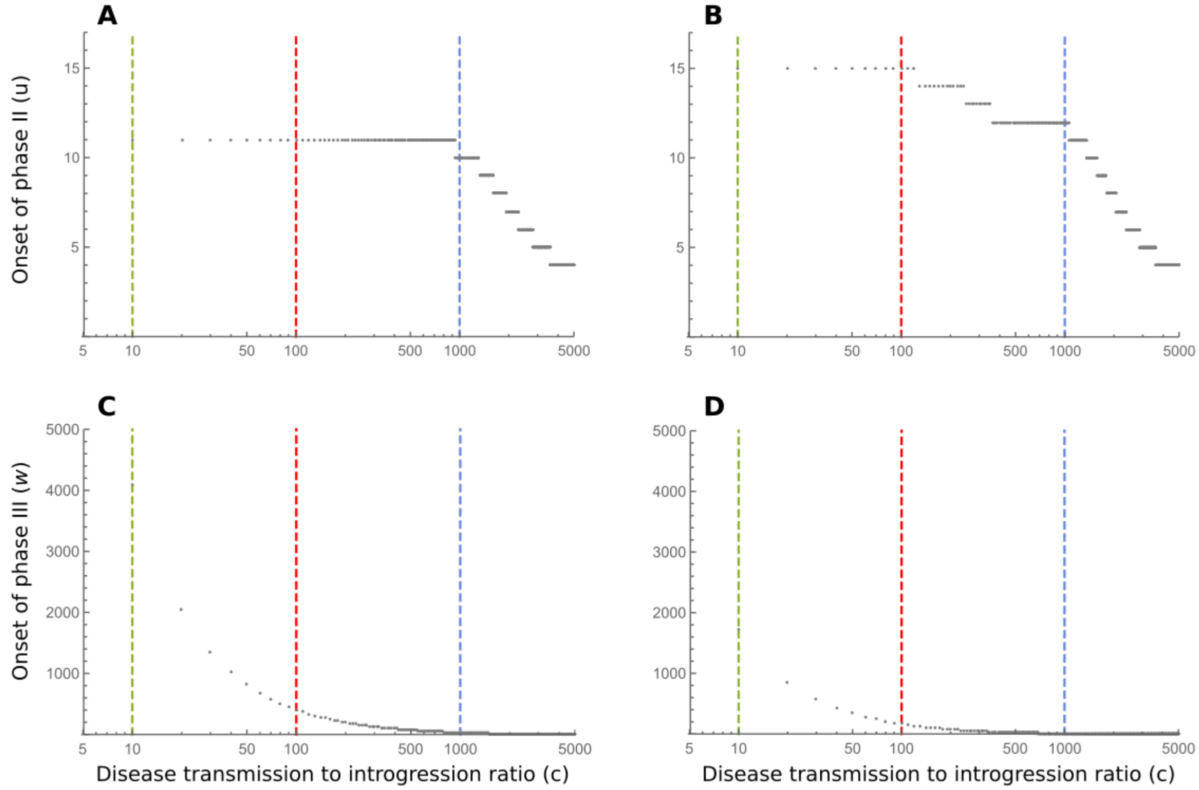


**Figure S3: Time of onset of phases II ($u$) and III ($w$) for different introgression to disease transmission ratios ($c$) in the two-time-scales model.** Dashed lines show the parameter values for which we show the dynamics explicitly, in the main text in Figures 3 and 4 (red), or in Fig. S4 (green and blue). (A) Onset of phase II for the symmetric case. (B) Onset of phase II for the asymmetric case ($P_1(0) = 60$, $P_2(0) = 100$). (C) Onset of phase III for the symmetric case. (D) Onset of phase III for the asymmetric case ($P_1(0) = 60$, $P_2(0) = 100$). Other than the the between-species contact rates, all other parameters are the same as those in Fig. 3 (for (A) and (C)) and Fig. 4 (for (B) and (D)). Notice that the $c$ parameter on the X-axis is plotted on a logarithmic scale.

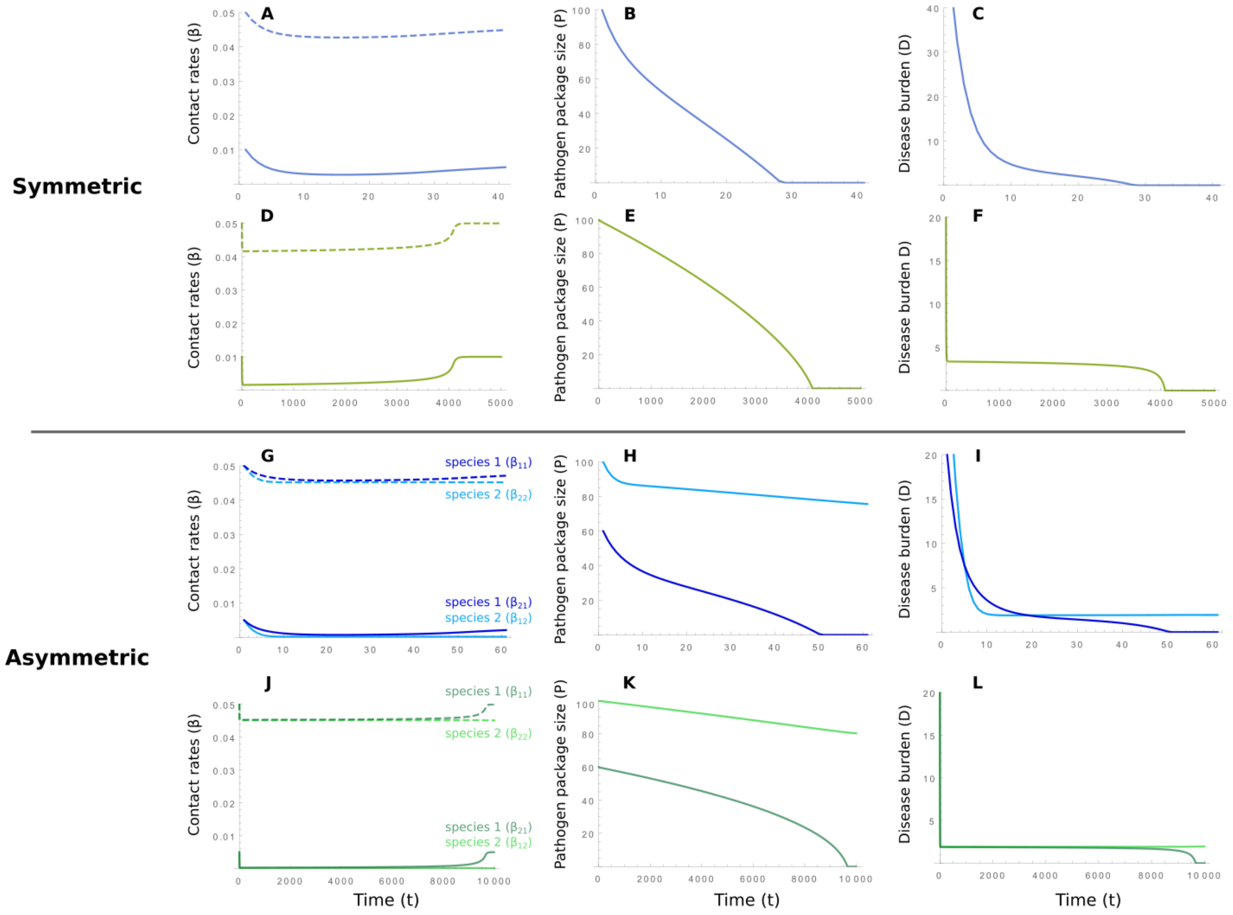


**Figure S4: Two examples of dynamics of the two-time-scales model for the introgression to disease transmission ratios ($c$) values marked in Fig. S3**. (A)–(F) show the results for the symmetric case, comparable with Fig. 3, and (G)–(L) show the results for the asymmetric case, comparable with Fig. 4. For the panels with blue curves (corresponding to the blue line in Fig. S3), disease transmission to introgression values are $c = 1000$, while the rest of the parameters are the same as those in Fig. 3 and Fig. 4. For the panels with green curves (corresponding to the green line in Fig. S3), $c$ values are $c = 10$, while the rest of the parameters are the same as those in Fig. 3 and Fig. 4. Notice that the times scales for the different panels are different.

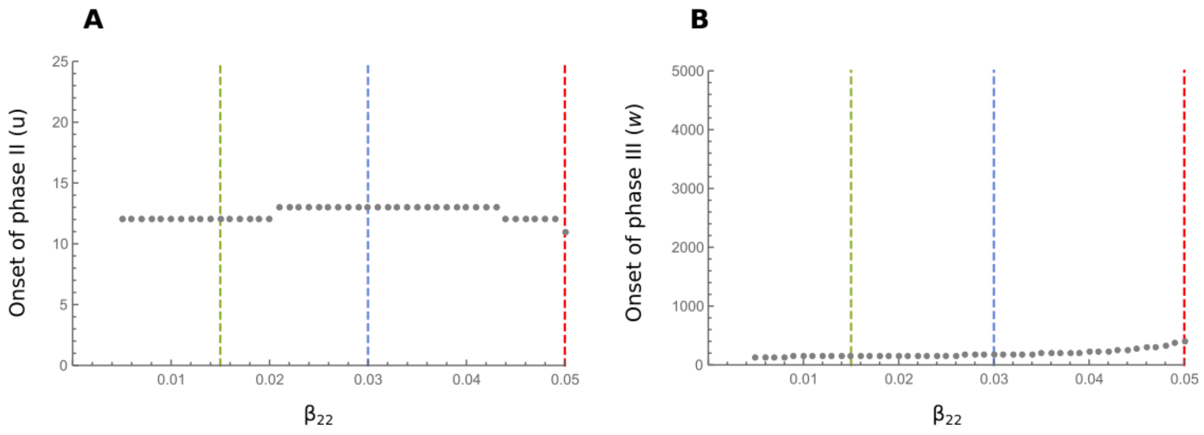


**Figure S5: Time of onset of phases II ($u$) and III ($w$) for different $\beta_{22}(0)$ values in the two-time-scales model.** Dashed lines show the parameter values for which we show the dynamics explicitly, in the main text in Figures 3 (red, which is the symmetric case of $\beta_{22}(0) = \beta_{11}(0) = 0.05$), or in Fig. S6 (green and blue). (A) Onset of phase II ($u$). (B) Onset of phase III ($w$).

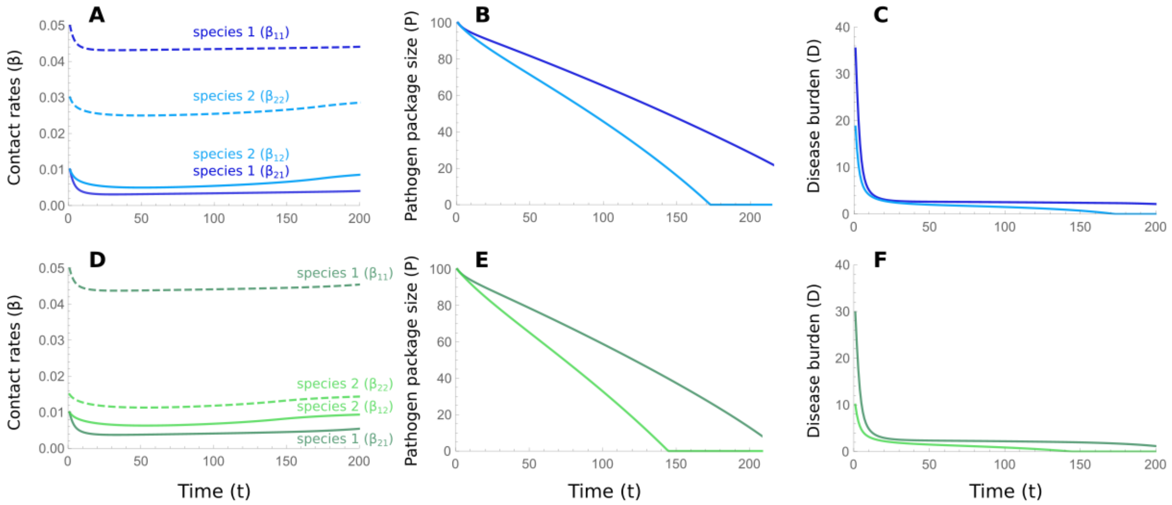


**Figure S6: Two examples of dynamics of the two-time-scales model for the $\beta_{22}(0)$ values marked in Fig. S5**. The results are comparable with Fig. 4. For the panels with blue curves (corresponding to the blue line in Fig. S5), within-species contact rates in species 2 are $\beta_{22}(0) = 0.03$, while the rest of the parameters are the same as those in Fig. 4. For the panels with green curves (corresponding to the green line in Fig. S5), $\beta_{22}(0)$ values are $\beta_{22}(0) = 0.015$, while the rest of the parameters are the same as those in Fig. 4.

#### A.2 Single-time-scale model

For the single-time-scale model, we conduct a similar analysis for exploring the parameter ranges to the one described above for the two-times-scales model. We vary one parameter at a time, using the scenarios in Figure 5 as our baseline. We evaluate the onset of phase II ($u$) and phase III ($w$), defined in a similar manner. Since this model is defined in continuous time, $u_{N_i}$ is defined as the earliest time in which the absolute value of the derivative of $N_i$ is smaller than $10^{-3}$. The onset of phase II, $u$, is defined as the maximum of $u_{N_1}$ and $u_{N_2}$. The onset of phase III, $w$, is defined as the earliest time, $t$, in which the disease burden is lifted in one of the species,i.e. either $\alpha_1(t) = 0$ or $\alpha_2(t) = 0$ ($w_{\alpha_1}$ and $w_{\alpha_2}$, respectively).

##### A.2.1 Between-species contact rates

We varied the between-species transmission rate, $\beta_a$, from 0.01 to 0.5 in increments of 0.01, for the symmetric and asymmetric cases. Figure S7 shows the onset of the phases for different $\beta_a$ values, and full dynamics for two examples ($\beta_a = 0.02$ in blue, and $\beta_a = 0.005$ in green) are shown in Figure S8. For the symmetric case, the onset of both phases decreases as the between-species contact rates increase, since faster contact rates translate to faster dynamics in the system. In the asymmetric case, the onset of phase III decreases as $\beta_a$ increases, but the onset of phase II initially increases but around $\beta_a = 0.4$ begins to increase as $\beta_a$ increases. This may be due to the complex feedback between introgression, disease transmission, and population dynamics, which may result in different dynamics at different parts of the parameter space.

The between-species contact rate $\beta_a$ plays a role both in determining inter-species transmission of disease (Eq. 8) and the rate of introgression (Eq. 9). In general, increase in $\beta_a$ leads to faster dynamics in which the onset of both phase II and phase III occur earlier; however, for very high $\beta_a$ and asymmetry in disease-related mortality rates, complex interaction in the equations lead to non-trivial changes in the onset of the phases (Fig. S7).

##### A.2.2 Ratio of introgression to disease transmission

We varied the introgression to disease transmission ratio from $c = 0.01$ to $c = 0.3$ in increments of 0.01. In the asymmetric case ($\alpha_1(0) = 0.5, \alpha_2(0) = 1$), with $c$ values above 0.3, $w$ becomes very low, eventually even lower than $u$. This is because we defined the onset of phase II, $u$, to be the maximum of $u_{N_1}$ and $u_{N_2}$, and $w$ to be the minimum of $w_{\alpha_1}$ and $w_{\alpha_2}$, and with very high rates of adaptive introgression one species may overcome disease burden before the other species has stabilized. Therefore, the notion of different phases, as discussed in the main text, does not apply to such high introgression rates. We therefore limited our range of investigation of this parameter to $c$ values up to 0.3.

Figure S9 shows the onset of the two phases for different $c$ values, and two examples of the dynamics ($c = 0.01$ and $c = 0.3$) are shown in Figure S10. The time of onset of phase II decreased as we increased $c$. The same is observed for the onset of phase III, due to higher $c$ resulting in faster adaptive introgression. These results are similar to those observed for the two-time-scales model (Fig. S3). For the symmetric case, the dynamics in the examples we focused on are qualitatively similar to those in Figure 5 (Fig. S10). For the asymmetric case, the dynamics have characteristics that are different than those in Figure S5, such as switches between whether $N_1 > N_2$ or $N_2 > N_1$, but some characteristics remain the same: high initial disease burden which rapidly decreases as population sizes decrease (phase I), followed by a relatively stable phase (phase II) with relatively constant low disease burden, followed by overcoming of disease burden of species 1 and increase in its population density (phase III).

The ratio between disease transmission and introgression, $c$, plays a role in Eq. 9, where higher $c$ values lead to faster introgression. Therefore, increasing $c$ results both in earlier onset of phase II and earlier onset of phase III (Fig. S9).

##### A.2.3 Initial asymmetry in population densities

As discussed above for the two-time-scales model, a plausible source of asymmetry, other than the characteristics of the pathogen package, is population densities. We therefore explored this source of asymmetry by considering the symmetric case in Figure 5 as our baseline, and varying the initial population density of species 2, $N_2(0)$ (initial population density of species 1 was kept as $N_1(0) = 2$). We kept the intrinsic demographic parameters, $K$, $\lambda$, and $\mu$ the same for both species. We varied the ratios between the two population densities from $N_2(0) = 0.01 \times N_1(0)$ to $N_2(0) = 1 \times N_1(0)$ in increments of 0.01.

Figure S11 shows the onset of the two phases for different ratios of initial population densities. Except for very high asymmetry, with species 1 being more than 20 times more dense than species 2, the onset of phase II decreased with more asymmetry. The onset of phase III decreased as the asymmetry was stronger. The examples in Figure S12 are qualitatively similar to those in Figure 5, with the three phases observable.

As with the two-time-scale model, for asymmetry in population density ($\beta_{22}(0)$ in the two-time-scales model, which could be interpreted as population density), the species that started out with the lower density, species 2, is the one that eventually overcomes disease burden faster, and consequently becomes denser than the other species.

The intial density of species 2, $N_2(0)$, plays a role in the initial conditions of

Eq. 7. When the initial population density in species 2 is decreased, it has less effect on species in terms of infections (Eq. 8). This results in species 1 initially maintaining higher population densities with lower $N_2(0)$ values (Fig. S12). This, in turns, results in higher rates of introgression in species 2 with lower $N_2(0)$, leading to earlier onset of phase III (Fig. S11). The onset of phase II is not simply determined with changes of $N_2(0)$.

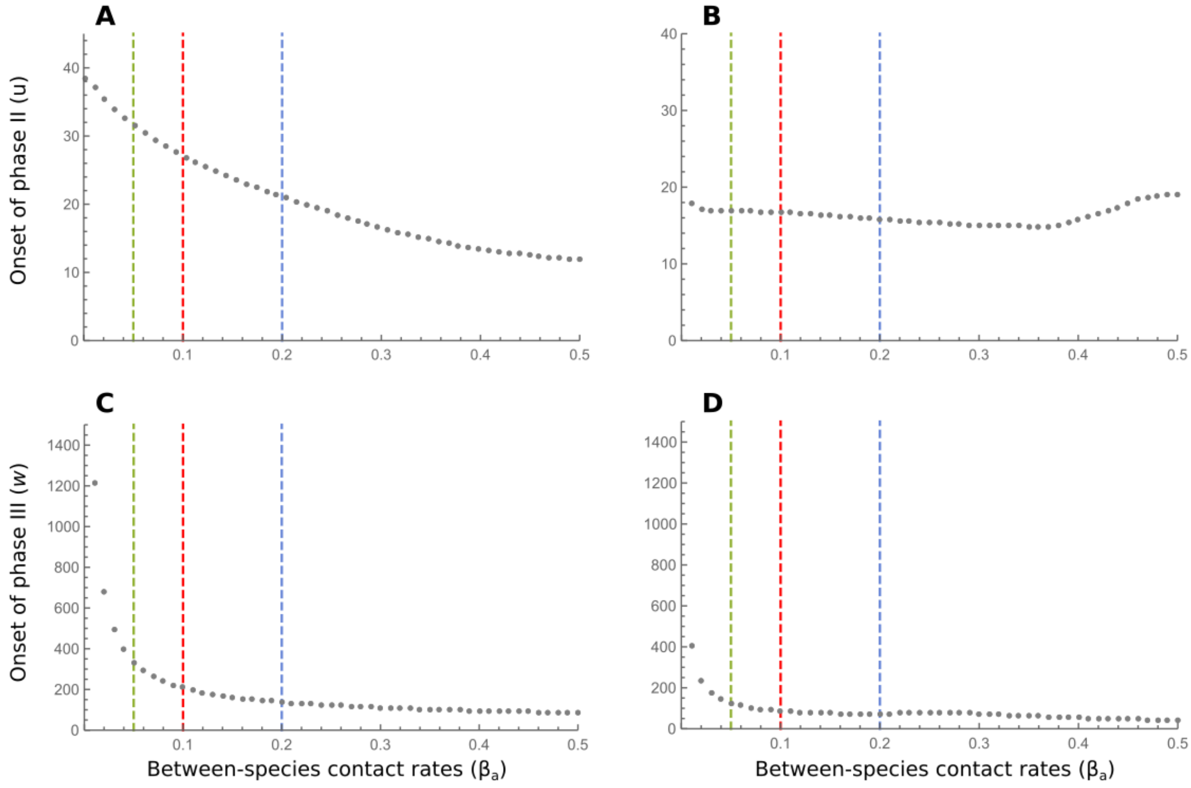


**Figure S7: Time of onset of phases II ($u$) and III ($w$) for different $\beta_a$ values in the single-time-scales model.** Dashed lines show the parameter values for which we show the dynamics explicitly, in the main text in Fig. 5 (red), or in Fig. S8 (green and blue). (A) Onset of phase II for the symmetric case. (B) Onset of phase II for the asymmetric case ($\alpha_1(0) = 0.5$, $\alpha_2(0) = 1$). (C) Onset of phase III for the symmetric case. (D) Onset of phase III for the asymmetric case ($\alpha_1(0) = 0.5$, $\alpha_2(0) = 1$). Other than the the parameter $\beta_a$, all other parameters are the same as those in Fig. 5.

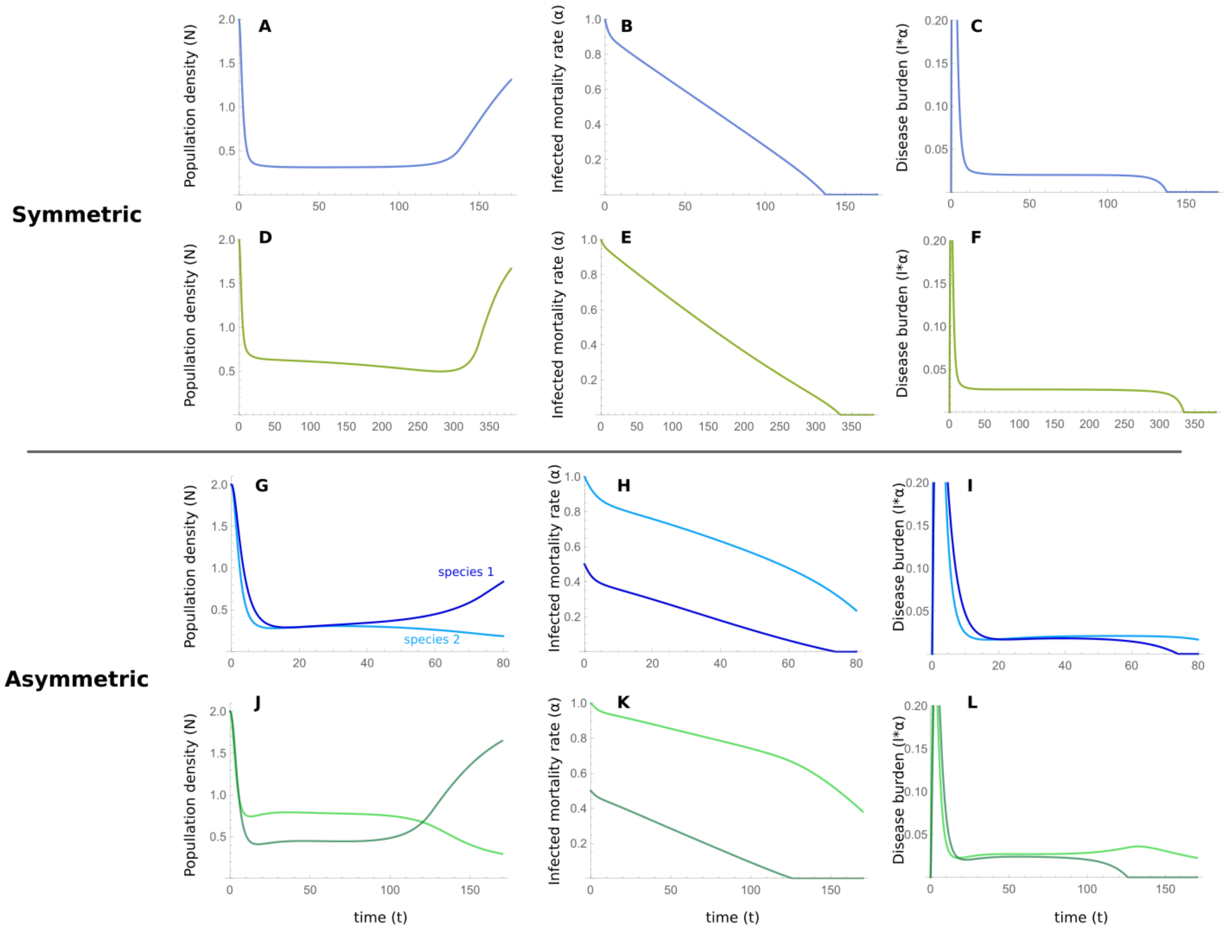


**Figure S8: Two examples of dynamics of the single-time-scale model for the $\beta_a$ values marked in Fig. S7**. (A)–(F) show the results for the symmetric case and (G)–(L) show the results for the asymmetric ($\alpha_1(0) = 0.5$, $\alpha_2(0) = 1$), comparable with Fig. 5. For the panels with blue curves (corresponding to the blue line in Fig. S7), between-species contact rates are $b_a = 0.2$, while the rest of the parameters are the same as those in Fig. 5. For the panels with green curves (corresponding to the green line in Fig. S7), between-species contact rates are $b_a = 0.05$, while the rest of the parameters are the same as those in Fig. 5. Notice that the times scales for the different panels are different.

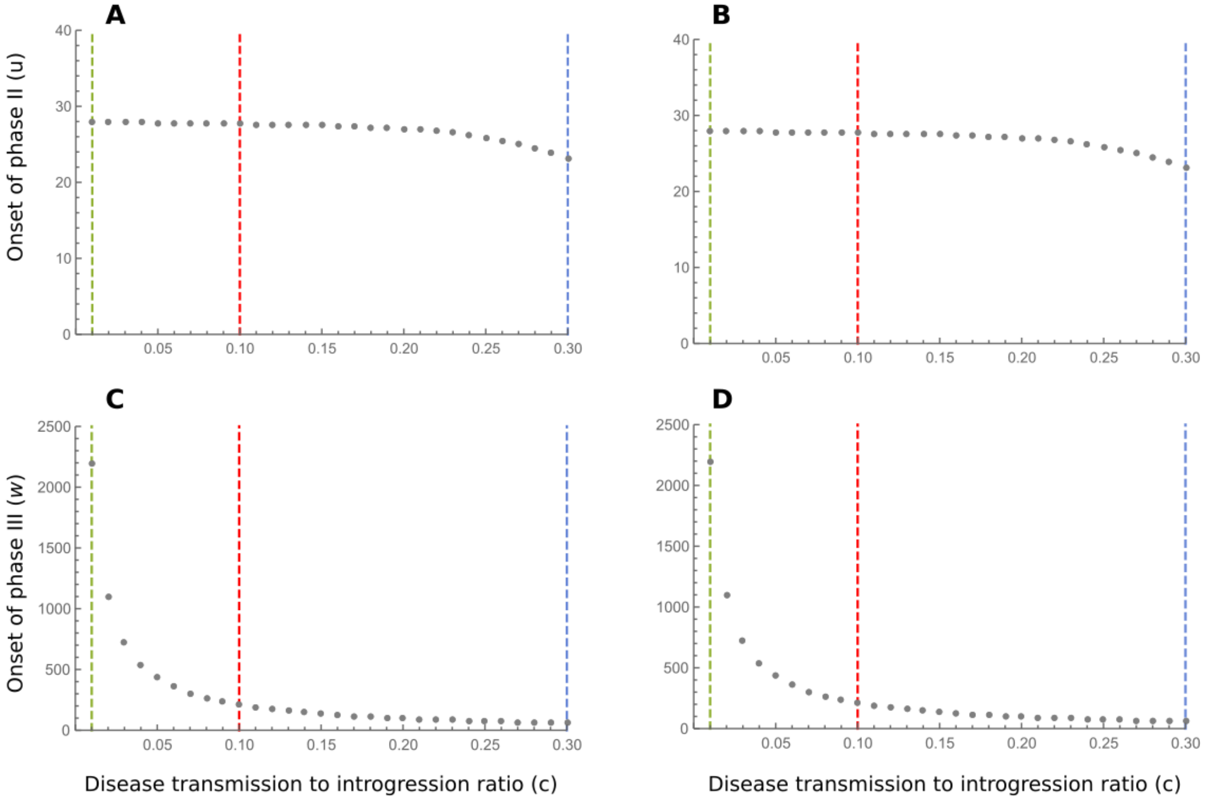


**Figure S9: Time of onset of phases II ($u$) and III ($w$) for different $c$ values in the single-time-scales model.** Dashed lines show the parameter values for which we show the dynamics explicitly, in the main text in Fig. 5 (red), or in Fig. S10 (green and blue). (A) Onset of phase II for the symmetric case. (B) Onset of phase II for the asymmetric case ($\alpha_1(0) = 0.5$, $\alpha_2(0) = 1$). (C) Onset of phase III for the symmetric case. (D) Onset of phase III for the asymmetric case ($\alpha_1(0) = 0.5$, $\alpha_2(0) = 1$). Other than the parameter $c$, all other parameters are the same as those in Fig. 5.

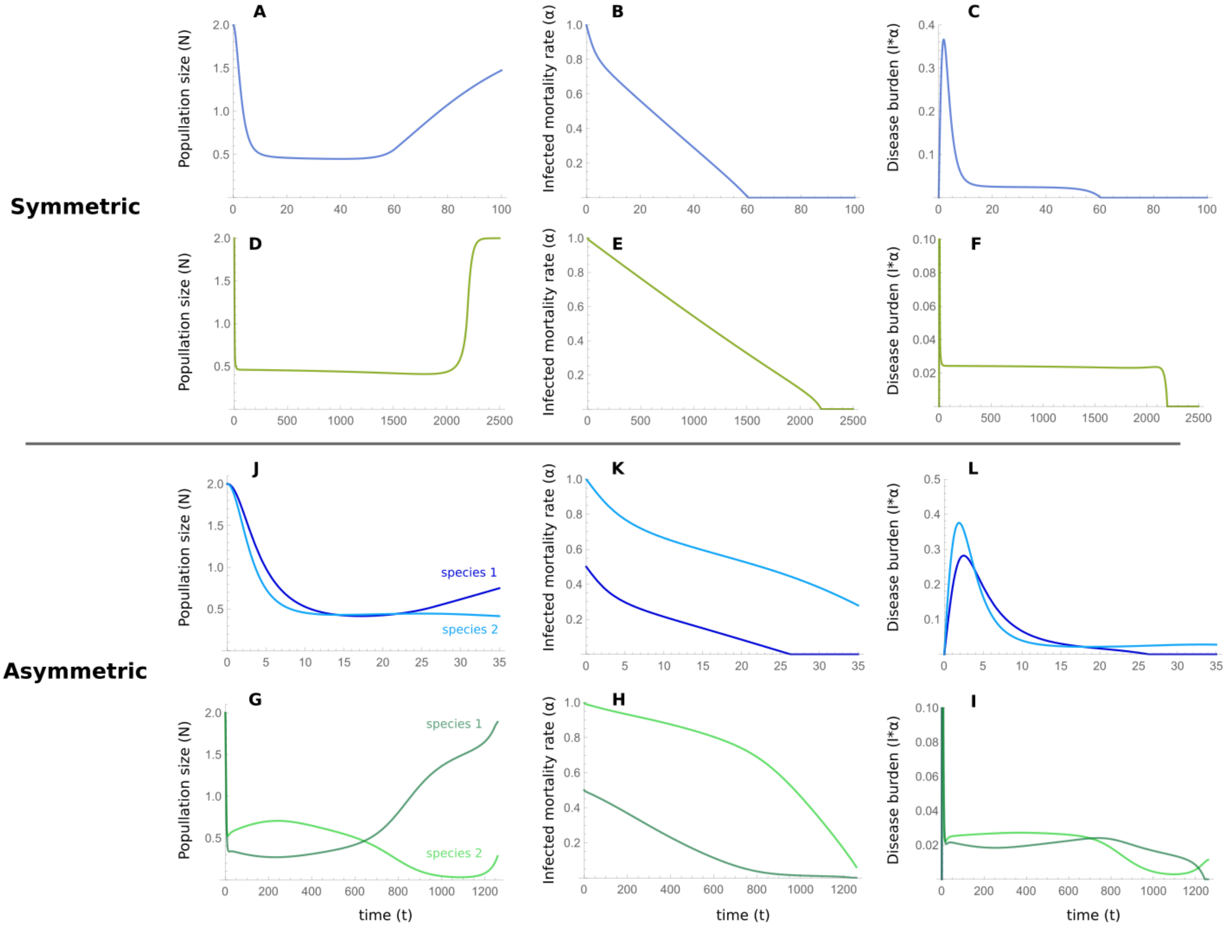


**Figure S10: Two examples of dynamics of the single-time-scale model for the $c$ values marked in Fig. S9**. (A)–(F) show the results for the symmetric case and (G)–(L) show the results for the asymmetric case ($\alpha_1(0) = 0.5$, $\alpha_2(0) = 1$), comparable with Fig. 5. For the panels with blue curves (corresponding to the blue line in Fig. S9), $c = 0.3$, while the rest of the parameters are the same as those in Fig. 5. For the panels with green curves (corresponding to the green line in Fig. S9), $c = 0.01$, while the rest of the parameters are the same as those in Fig. 5. Notice that the times scales for the different panels are different.

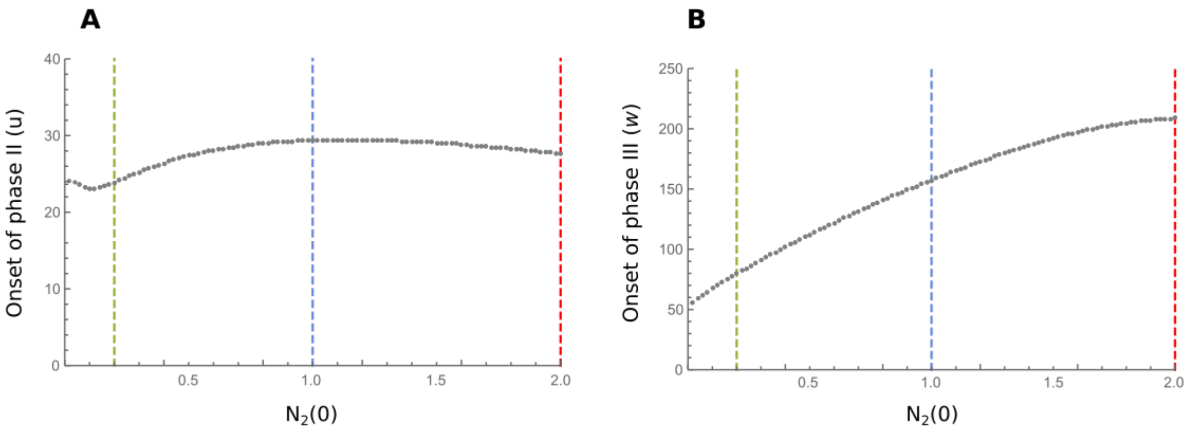


**Figure S11: Time of onset of phases II ($u$) and III ($w$) for different $N_2(0)$ values in the single-time-scales model.** Dashed lines show the parameter values for which we show the dynamics explicitly, in the main text in Figures 5 (red), or in Fig. S12 (green and blue). (A) Onset of phase II ($u$). (B) Onset of phase III ($w$).

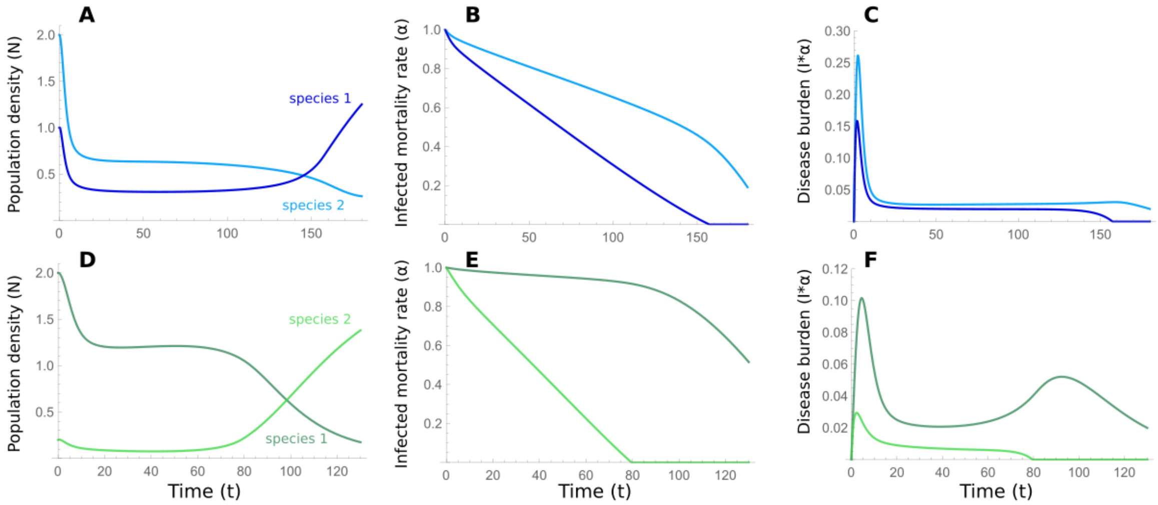


**Figure S12: Two examples of dynamics of the single-time-scale model for the $N_2(0)$ values marked in Fig. S11**. The results are comparable with Fig. 5. For the panels with blue curves (corresponding to the blue line in Fig. S11), $N_2(0) = 0.5$, while the rest of the parameters are the same as those in Fig. 5. For the panels with green curves (corresponding to the green line in Fig. S11), $N_2(0) = 0.1$, while the rest of the parameters are the same as those in Fig. 5. Notice that the times scales for the different panels are different.

### B Modeling regular adaptation in addition to adaptive introgression

In this section we explore adding adaptation to disease due to de novo immune-related mutations in both species, in addition to introgression.

#### B.1 Two-time-scales model

In the two-time-scales model, adaptation to the novel pathogen package is measured in terms of the number of pathogens to which each of the species is vulnerable. While introgression depends on inter-species contact rates, regular within-species adaptation is expected to be independent of these. In many cases, adaptations are acquired by natural selection acting on newly arisen mutations or on standing variation. The process by which this occurs depends on several factors, including the strength of selection, the effective population size and genetic drift, and the rate of mutation.

Modeling the population genetics of this process in the context of our model is not a simple task. We therefore decided to make many simplifying assumption, and instead of incorporating a population genetic model, we assume that beneficial immune-related mutations, which reduce the experienced pathogen package size $P$, appear and are driven to fixation at a constant rate (see [1–6] for similarly simplified models). Moreover, although fixation through selection is not an instantaneous process, and in intermediate steps the population would be composed of both individuals carrying the immune-related alleles and those that do not carry them, we assume that the time between appearance and fixation is negligible at the evolutionary time scale. These assumptions, which may not be appropriate in many cases, allow us to model regular adaptation as a single parameter, $\mu_a$, which is the (constant) rate at which immune-related mutations that are eventually driven to fixation in the population convey tolerance to a single pathogen. This rate is measured according to the evolutionary time-scale of this model ($t$). Therefore, at each time step $t$, species $i$ evolves tolerance to $\mu_a P_i(t)$ novel pathogens (with the assumption that the rate of evolution of tolerance to each pathogen is the same, and that the evolutionary processes for each pathogen are independent). We also assume that these rates are the same for both species.

The equation describing the pathogen package size is a modified version of Eq. 5, namely :

$$\begin{aligned} P_1(t) &= \max\left\{(1-\mu_a)P_1(t-1) - c\,\beta_{21}(t-1), 0\right\} \\ P_2(t) &= \max\left\{(1-\mu_a)P_2(t-1) - c\,\beta_{12}(t-1), 0\right\}, \end{aligned} \tag{S1}$$

and the two-time-scales model with regular adaptation is identical to the one described in the main text but with Eq. S1 replacing Eq. 5.

For this model, we varied the $\mu_a$ parameter from 0 to 0.05 in increments of 0.001, with the other parameters remaining fixed to the baseline values of Fig. 3. We used the same definition of time of onset of phases II and III as presented in Appendix A to explore the effect of regular adaptation on the phases (Fig. S13). We also show two examples of the dynamics ($\mu_a = 0.005$ in green and $\mu_a = 0.01$ in blue) in Figure S14.

The examples in Figure S14 demonstrate that the dynamics are qualitatively similar to those without regular adaptation, for the parameters we investigated, in the sense that the three phases can be observed. In Figure S13, it can be seen that the onset of phase II is almost unchanged with increased regular adaptation, with a slight decrease, but the onset of phase III is earlier as the level of regular adaptation increases. This is so since in our model, we added regular adaptation in addition to adaptive introgression, in essence modeling two pathways by which tolerance can be evolved, and therefore the species overcome disease burden sooner than with adaptive introgression alone.

#### B.2 Single-time-scale model

We also incorporated regular adaptation in the single time-scale model, with the same simplifying assumption and the same reservations. Here, we introduce a parameter $\mu_a$, which is the constant rate of appearance of mutations that convey tolerance to pathogens, and which are eventually driven to fixation. Since we model population densities explicitly in this model, and the rate of appearance of novel mutations in the populations is proportional to population size, we model the effect of regular adaptation as a reduction of disease-induced mortality, $\alpha_i$, by the rate of *de novo* appearance of adaptive mutations in the population, $N_i\mu_b$:

$$\begin{aligned}
\frac{d\alpha_1(t)}{dt} &= -c\beta_a N_2(t) - \mu_b N_1(t) \quad \text{provided } \alpha_1(t) > 0\text{, otherwise } \frac{d\alpha_1(t)}{dt} = 0 \\
\frac{d\alpha_2(t)}{dt} &= -c\beta_a N_1(t) - \mu_b N_2(t) \quad \text{provided } \alpha_2(t) > 0\text{, otherwise } \frac{d\alpha_2(t)}{dt} = 0 \qquad \text{(S2)}
\end{aligned}$$

The single-time-scale model with regular adaptation is the same as in the main text except that Eq. S2 replaces Eq. 9.

We varied the $\mu_b$ parameter from 0 to 0.05 in increments of 0.001, with the other parameters remaining fixed to the baseline values of Fig. 5 (symmetric case). We used the same definition of time of onset of phases II and III as presented in Appendix A to explore the effect of regular adaptation on the phases (Fig. S15). We also show two examples of the dynamics ($\mu_a = 0.005$ in green and $\mu = 0.01$ in blue) in Figure S16.

As with the two-time-scales model, the examples for different rates of $\mu_b$ show similar dynamics with the three phases described in the main text (Fig. S16). For higher rates of regular adaptation, the onset of the second phase does not change much in the parameter range range we investigated, but the onset of phase II is significantly reduced (Fig. S15). As was discussed for the two-time-scales model, we model regular adaptation as a process which occurs in parallel and in addition to adaptive introgression, and therefore disease burden is overcome more rapidly with more regular adaptation.

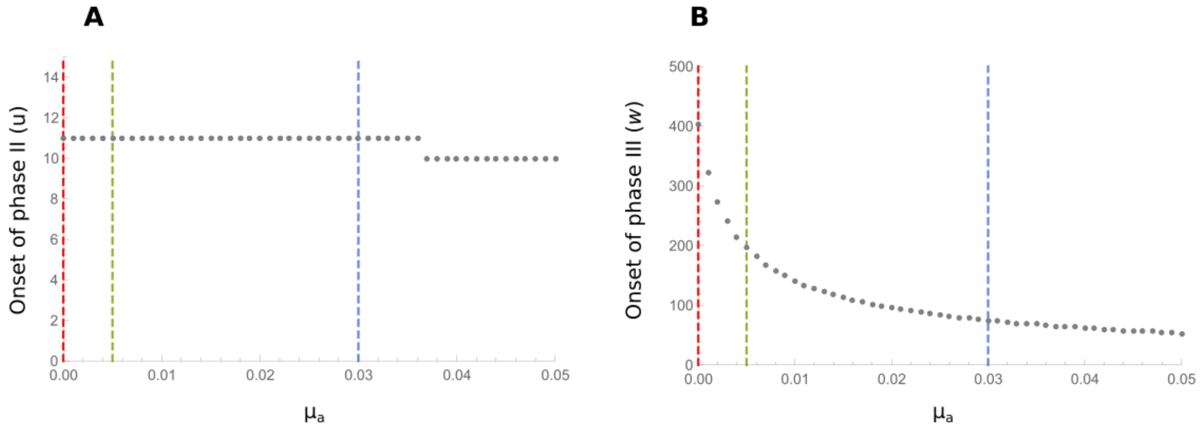


**Figure S13: Time of onset of phases II ($u$) and III ($w$) for different $\mu_a$ values in the two-time-scales model with regular adaptation.** Dashed lines show the parameter values for which we show the dynamics explicitly, in the main text in Figures 3 (red), or in Fig. S14 (green and blue). (A) Onset of phase II ($u$). (B) Onset of phase III ($w$).

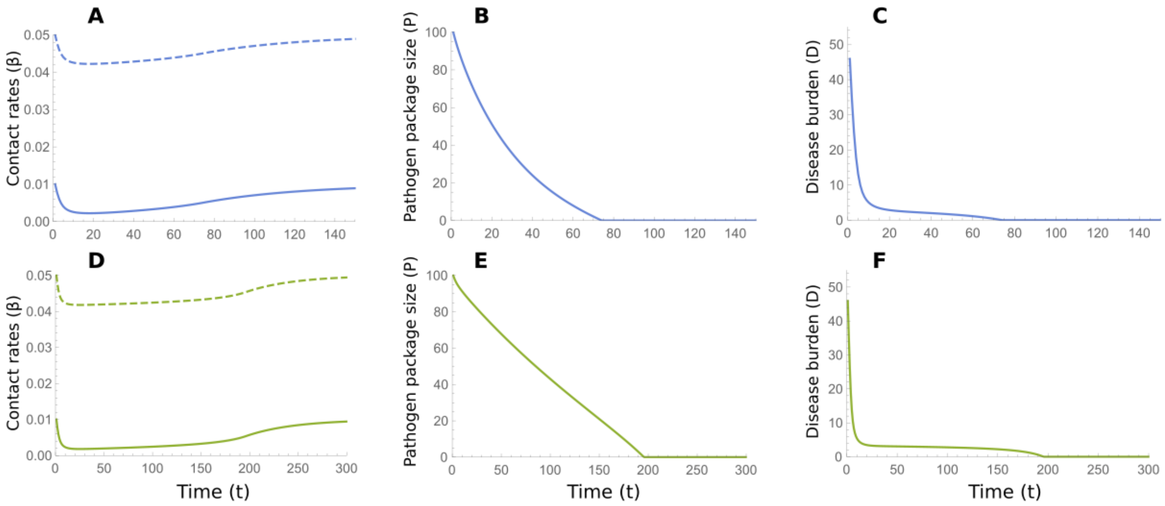


**Figure S14: Two examples of dynamics of the two-time-scales model with regular adaptation for the values marked in Fig. S13**. For the panels with blue curves (corresponding to the blue line in Fig. S13), $\mu_a = 0.03$, while the rest of the parameters are the same as those in Fig. 3. For the panels with green curves (corresponding to the green line in Fig. S13), $\mu_a = 0.0005$, while the rest of the parameters are the same as those in Fig. 3. Notice that the times scales for the different panels are different.

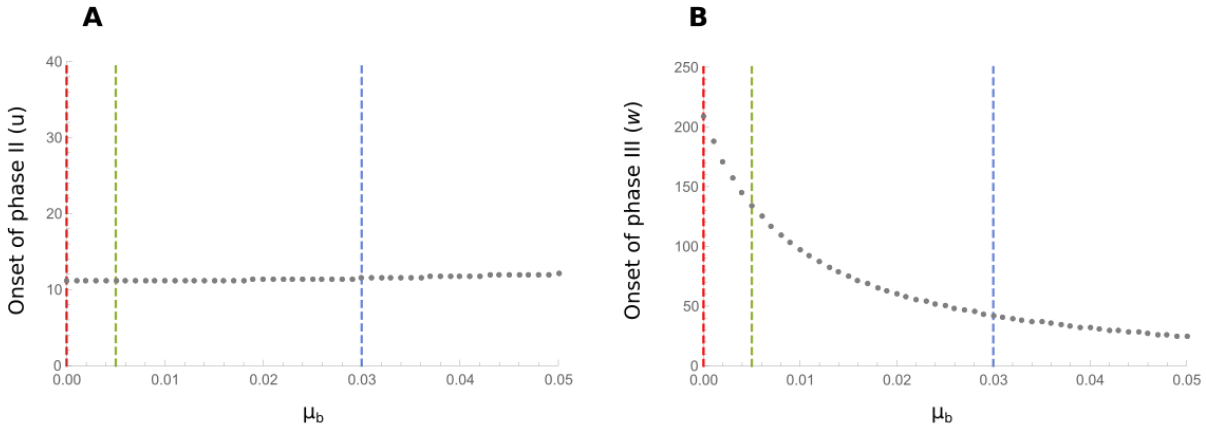


**Figure S15: Time of onset of phases II ($u$) and III ($w$) for different $\mu_b$ values in the single-time-scale model with regular adaptation.** Dashed lines show the parameter values for which we show the dynamics explicitly, in the main text in Figures 5 (red), or in Fig. S16 (green and blue). (A) Onset of phase II ($u$). (B) Onset of phase III ($w$).

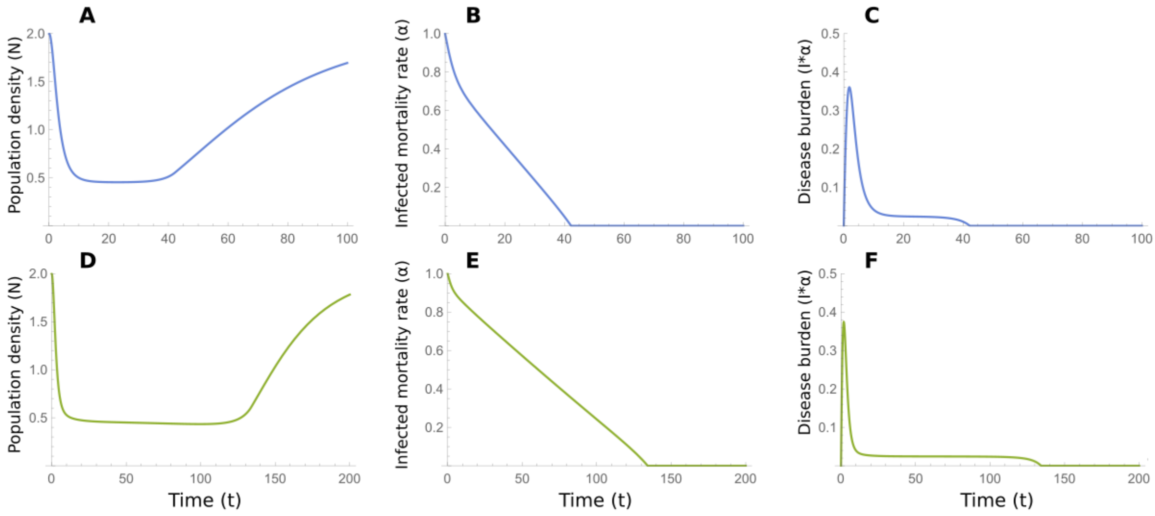


**Figure S16: Two examples of dynamics of the single-time-scale model with regular adaptation for the values marked in Fig. S15**. For the panels with blue curves (corresponding to the blue line in Fig. S15), $\mu_a = 0.03$, while the rest of the parameters are the same as those in Fig. 5 (symmetric case). For the panels with green curves (corresponding to the green line in Fig. S15), $\mu_a = 0.0005$, while the rest of the parameters are the same as those in Fig. 5 (symmetric case). Notice that the times scales for the different panels are different.

### C Modeling inter-species competition

Given the physical similarity and close phylogenetic relations of Neanderthals and Moderns, it is plausible that when they interacted in the Levant they were in competition for similar resources, and that they occupied overlapping ecological niches [7, 8]. In the models presented in the main text, we did not incorporate direct competition between the species. In the two-time-scales model, where demography is only implicitly modeled, incorporating competition would present some difficulty. However, in the single-time-scale model, demographics are explicitly modeled, and therefore, it is easier to accommodate competition between the species in the model, and explore how that might affect the dynamics we describe. In this section we relax the assumption of no competition in the single-time-scale model, and incorporate the effect of competition on growth rates.

For this purpose, we introduce a competition parameter $\phi$, which measures the niche overlap of the two species, i.e. for $\phi = 0$, the species' niches do not overlap and there is no competition, and for $\phi = 1$, the niches are identical and inter-species competition is the same as within-species competition [9]. In the main text, we modeled density-dependent growth rate as a function of the intrinsic growth rate $\lambda$ and the half-maximum population density $K$, and now we incorporate inter-species competition:

$$\Lambda_i(N_i, N_j) = \frac{\lambda N_i}{K + (N_i + \phi N_j)}, \quad i \neq j, \; i, j = 1, 2. \tag{S3}$$

The single-time-scale model with competition is, therefore, the model described in the main text with Eq. S3 used instead of Eq. 6 (for $\phi = 0$, this model converges to the single-time-scale model presented in the main text).

With increased competition between the species, the onset of phase II is slightly delayed, and the onset of phase III is more substantially delayed (Fig. S17). Therefore, competition would act to increase the duration of the second stable phase, for the parameters we explored (Fig. S18). For the two examples we investigated, the dynamics are qualitatively similar to those presented in the main text, as the three phases can be observed.

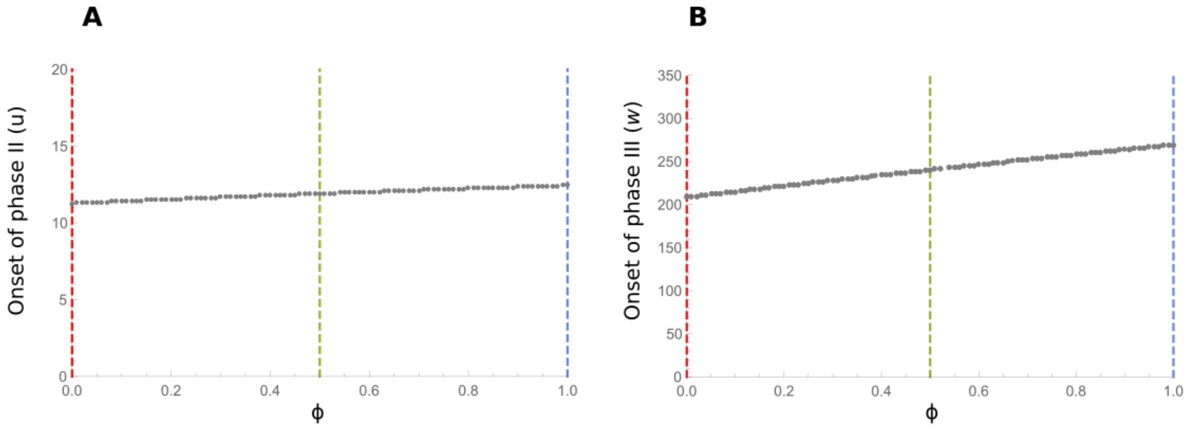


**Figure S17: Time of onset of phases II ($u$) and III ($w$) for different $\phi$ values in the single-time-scale model with competition.** Dashed lines show the parameter values for which we show the dynamics explicitly, in the main text in Figures 5 (red), or in Fig. S17 (green and blue). (A) onset of phase II ($u$). (B) Onset of phase III ($w$).

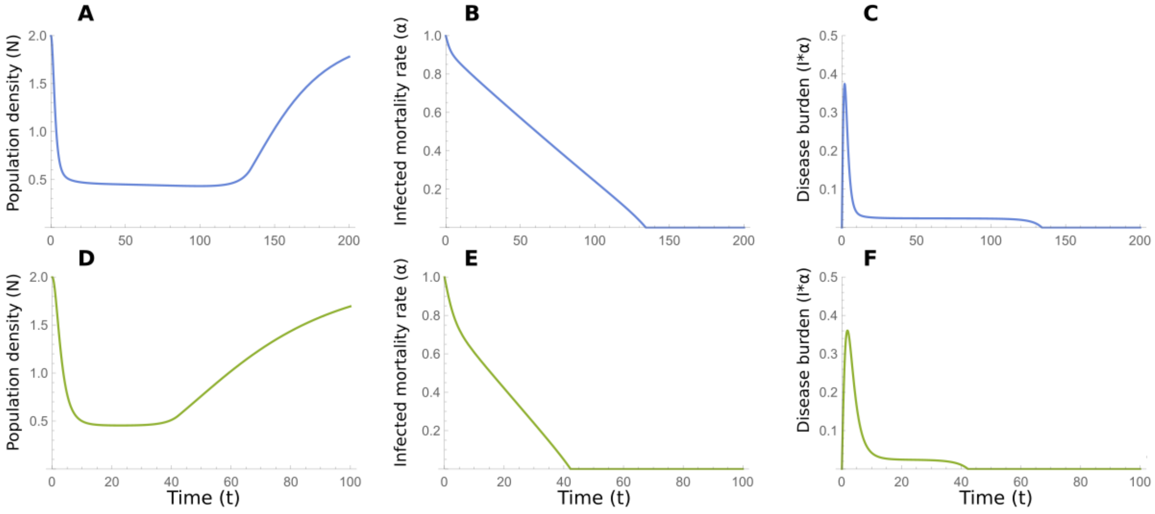


**Figure S18: Two examples of dynamics of the single-time-scale model with competition for the values marked in Fig. S17**. For the panels with blue curves (corresponding to the blue line in Fig. S17), $\phi = 1$ (full niche overlap), while the rest of the parameters are the same as those in Fig. 5 (symmetric case). For the panels with green curves (corresponding to the green line in Fig. S17), $\phi = 0.5$ (partial niche overlap), while the rest of the parameters are the same as those in Fig. 5 (symmetric case). Notice that the times scales for the different panels are different.
